# Conserved interfaces mediate multiple protein-protein interactions in a prokaryotic metabolon

**DOI:** 10.1101/2024.02.17.580827

**Authors:** Sanchari Bhattacharyya, Srivastav Ranganathan, Sourav Chowdhury, Bharat V Adkar, Mark Khrapko, Eugene I Shakhnovich

## Abstract

Enzymes in a pathway often form metabolons through weak protein-protein interactions (PPI) that localize and protect labile metabolites. Due to their transient nature, the structural architecture of these enzyme assemblies has largely remained elusive, limiting our abilities to re-engineer novel metabolic pathways. Here we delineate a complete PPI map of 1225 interactions in the *E. coli* 1-carbon metabolism pathway using bimolecular fluorescence complementation that can capture transient interactions *in vivo* and show strong intra- and inter-pathway clusters within the folate and purine biosynthesis pathways. Scanning mutagenesis experiments along with AlphaFold predictions and meta-dynamics simulations reveal that most proteins use conserved “dedicated” interfaces distant from their active sites to interact with multiple partners. Diffusion-reaction simulations with shared interaction surfaces and realistic PPI networks reveal a dramatic speedup in metabolic pathway fluxes. Overall, this study sheds light on the fundamental features of metabolon biophysics and structural aspects of transient binary complexes.

**Significance statement:** Enzymes from the same metabolic pathway often form dynamic assemblies called metabolons, which channel metabolites as well as protect labile intermediates. Yet very little is known about their structural features or what makes these interactions transient. Paucity of such information has particularly affected our ability to engineer novel metabolic pathways, construct multi-scale mathematical models of cells, etc. We address this by obtaining a comprehensive map for 1225 interaction pairs in the essential 1-carbon metabolism pathway of *E. coli*. Using both high-throughput experiments and computation, we uncover that metabolon proteins tend to use a conserved dedicated interface to interact with their partners. These results shed light on structural and energetic aspects of PPI in metabolons at near atomic level of resolution.

## Introduction

Unlike purified enzymes in a test tube, enzymes inside cells often do not function in isolation. In many cases, they carry out their reaction through formation of multi-enzyme assemblies. Metabolons - a term first coined by Paul Srere - are one such protein complex, which are transiently interacting supramolecular entities formed through weak interactions among sequential enzymes in a metabolic pathway which help to channel metabolites (1, 2). In higher eukaryotes, there has been evidence for such complexes that dynamically associate and dissociate in response to cellular stimuli, the most notable of them being purinosomes (3–5). It is generally postulated that formation of a metabolon has several advantages like metabolite channeling, increasing local concentration of metabolites, preventing competing pathways and protection of unstable and cyto-toxic intermediates (6, 7). Despite this, mechanistic understanding of how proteins in a metabolon interact have remained elusive. Moreover, due to the transient nature of the interactions, traditional methods of structural biology have failed to address these questions. These problems have largely impaired our abilities to design or redesign new enzyme pathways in metabolic engineering. Based on information from cross-linking studies and mass spectrometry, Wu and Minteer (8) built a low-resolution structure of the tri-carboxylic acid cycle (TCA) metabolon using constraint based protein-protein docking. This study gave the first structural insight into a metabolon and showed evidence of electrostatic channeling upon association of proteins (8).

More recently, it has been shown that even prokaryotic cells, which were once thought to be a bag of freely floating enzymes and metabolites, show evidence of metabolon formation and are therefore able to achieve a substantial degree of metabolic compartmentalization. In our previous work (9), using immunoprecipitation experiments, we have shown that Dihydrofolate Reductase (DHFR), a central enzyme of the *E. coli* 1-carbon metabolism pathway, associates with multiple proteins in its functional vicinity via weak interactions, which in turn was also responsible for overexpression toxicity of DHFR. Though precise evidence of substrate channeling was not established, a more recent work (10) presented compelling circumstantial evidence that much of these interactions involve channeling of substrates, causing diffusion limitation of the substrates inside the cell. To obtain a deeper understanding of the breadth and nature of interactions in prokaryotic enzyme assemblies, in this work we established a complete PPI map of the *E. coli* 1-carbon metabolism pathway using biomolecular fluorescence complementation assay. Further, using high throughput library generation, screening, along with computational simulations using AlphaFold and metadynamics, we answer several key questions about the structural architecture of such transient enzyme assemblies. We discover that most proteins utilize conserved structural interfaces dedicated to interacting with multiple partners, thereby behaving as date hubs as opposed to more permanent multi-enzyme assemblies or party hubs. Finally, using a coarse-grained diffusion-reaction model, we show that such enzyme interactions cause a dramatic enhancement of reaction fluxes that are several orders of magnitude greater as compared to a scenario with no PPI. This study not only represents the first detailed investigation of PPIs forming metabolons in prokaryotic enzyme pathways but also represents the first high-resolution structural insight into the precise molecular architecture of such fleeting miniature protein assemblies.

## Results

### Split YFP system to measure interactions in the 1-carbon metabolism pathway

The 1-carbon metabolism pathway is an essential pathway consisting of 3 inter-dependent biosynthetic pathways that catalyze the transfer of 1-carbon units from methyl donors (5,10-methelene THF, 5-methyl THF, N-formyl THF) to generate purine and pyrimidine nucleotides as well as methionine and pantothenic acid. The methyl donors themselves are produced through the folate biosynthesis pathway that starts with GTP and chorismate and involves condensation of para-amino benzoate (PABA) with 6-hydroxymethyl-dihydropterin diphosphate to form 7,8-dihydropteroate, which is eventually converted to dihydrofolate (DHF) and tetrahydrofolate (THF) (Fig 1A). The pyrimidine biosynthesis pathway intersects the folate pathway at its second last step catalyzed by thymidylate synthase (ThyA), where uridine monophosphate (dUMP) accepts a methyl group from 5,10-methylene THF to convert to thymidine monophosphate (dTMP). The purine biosynthesis pathway utilizes folate derivatives twice: in the third step where the enzyme PurT catalyzes conversion of Glycineamide ribonucleotide (GAR) to formyl-GAR with the help of formate; in the penultimate step where PurH catalyzes the conversion of AICAR to formyl-AICAR using N-formyl THF as a cofactor.

**Figure 1:**
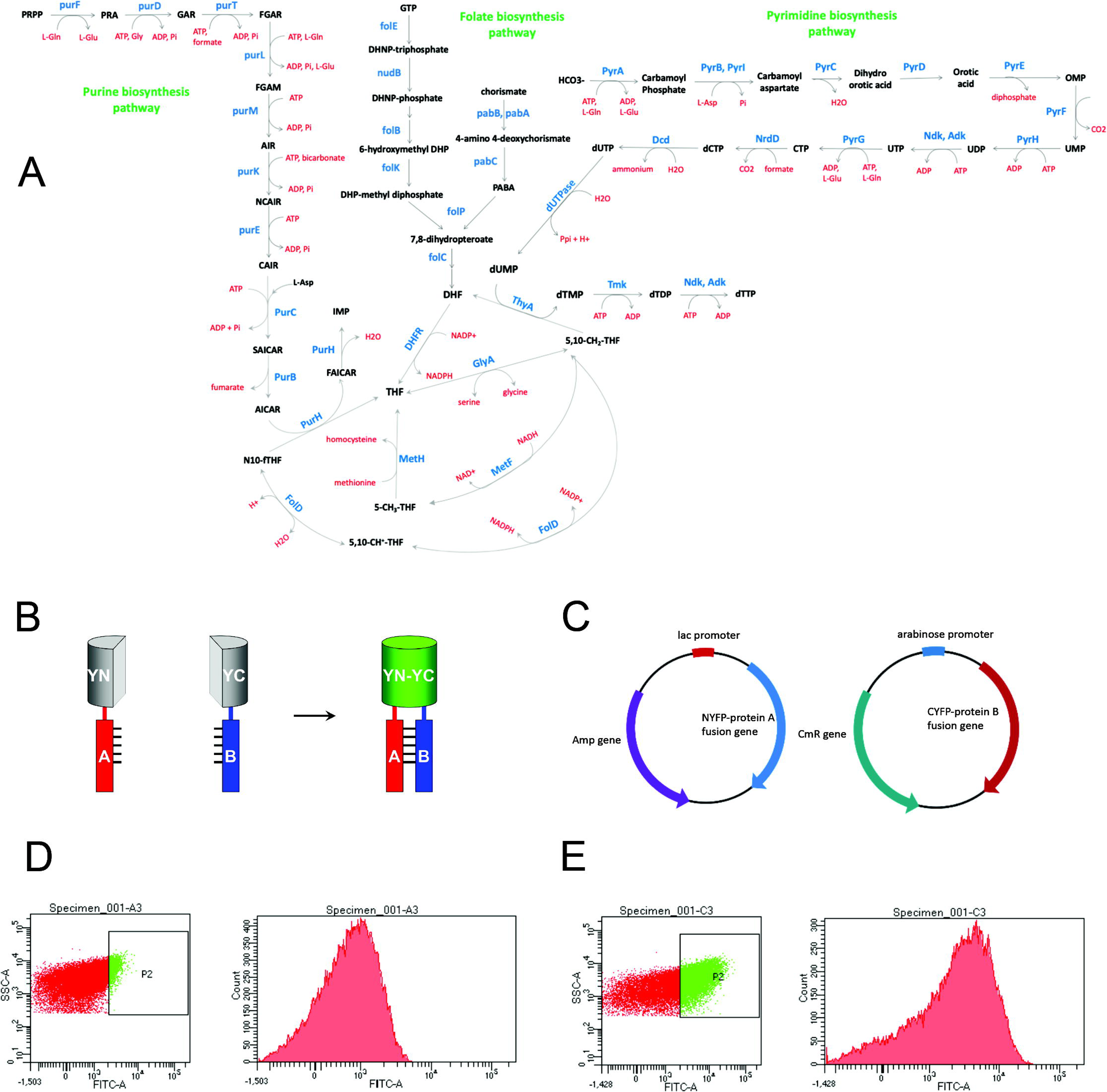
Overview of the E. coli 1-carbon metabolism pathway and YFP complementation method used to detect protein-protein interaction (PPI). (A) Purine, pyrimidine and folate biosynthesis pathways converge on to the 1-carbon metabolic pathway. (B) PPI is detected by bimolecular fluorescence complementation assay (BiFc) using the split YFP system wherein N-terminus of YFP molecule is fused to protein A, while the C-terminus is fused to protein B. A mature fluorescent YFP molecule is formed by complementation only if protein A and protein B interact. (C) A mutually compatible two-plasmid based system used for expression of YFP-fusion proteins. NYFP-protein A is expressed from an IPTG-inducible lac promoter, while CYFP-protein B is expressed from an arabinose inducible pBAD promoter. Representative flow-cytometry data for (D) a weakly interacting PPI pair and (E) strongly interacting pair. For both panels D and E, the left panel shows the SSC (side scatter of cells) as a function of fluorescence. SSC is a measure of internal granularity of the cell, which helps to identify a particular cellular population. P2 highlights the population that shows higher fluorescence than untransformed *E.coli* cells (not shown). This population is for visualization purpose only and is not used to calculate any relevant parameter for this study. The histogram on the right panel is derived from all cells (red and green dots together) in the left panel, and the mean fluorescence intensity (MFI) of this histogram is used to assess the extent of interaction.

Our previous work (9) had shown that enzymes in the folate biosynthesis pathway not only interact among themselves but also with those from functionally related purine and pyrimidine biosynthesis pathways. To systematically map interactions among all proteins in these three related pathways, we used a high-throughput way to measure these interactions. We used bimolecular fluorescence complementation assay (BiFc) using the split YFP system to measure interaction between pairs of proteins. In this system, one protein is fused to the N-terminus of YFP, while the other protein is fused to the C-terminus. A mature YFP is formed only if proteins A and B interact (Fig 1B). The two proteins are expressed from two mutually compatible plasmids and using two different inducers (IPTG based, and Arabinose based) (Fig 1C). Stronger interactions give higher fluorescence (Fig 1E), while weak or non-specific interactions give low fluorescence (Fig 1D).

### Matrix of 1225 interactions show large variation in PPI strengths

Out of total 41 proteins in the three pathways, genes corresponding to 35 proteins were cloned successfully as both N- and C-YFP fusions, which were used to generate a 35×35 matrix of 1225 interactions. For each pair, fluorescence was measured in two configurations: NYFP-X/CYFP-Y and NYFP-Y/CYFP-X, where X and Y are the two proteins in question. The entire dataset with interactions in two configurations in shown in Fig EV1A and Table EV2. In many cases, we observed strong interaction only in one configuration (either NYFP-X/CYFP-Y and NYFP- Y/CYFP-X), and not in both. This is presumably because fusion in one configuration may not be optimal for interaction. We therefore generated a symmetric matrix (Fig 2A and Table EV1) by selecting the highest fluorescence value obtained in any of the configuration. For a few pairs, the bacterial cultures co-transformed with both plasmids grew very poorly, hence significant number of cells could not be analyzed (grey boxes). For the remaining pairs, mean fluorescence intensity (MFI) showed a wide range of variations (Fig EV2A). As seen from Fig 2A, vast majority of the interactions are weak (MFI < 750). Several diagonal elements in the table show strong interactions, which correspond to formation of homo-oligomers. Interestingly, several off-diagonal protein pairs showed highly discernible interaction (MFI > 750). We classified them into weak (blue for 750 < MFI < 900), moderate (yellow for 900 < MFI < 1100) and strong (red boxes indicate MFI > 1100) interactions.

**Figure 2:**
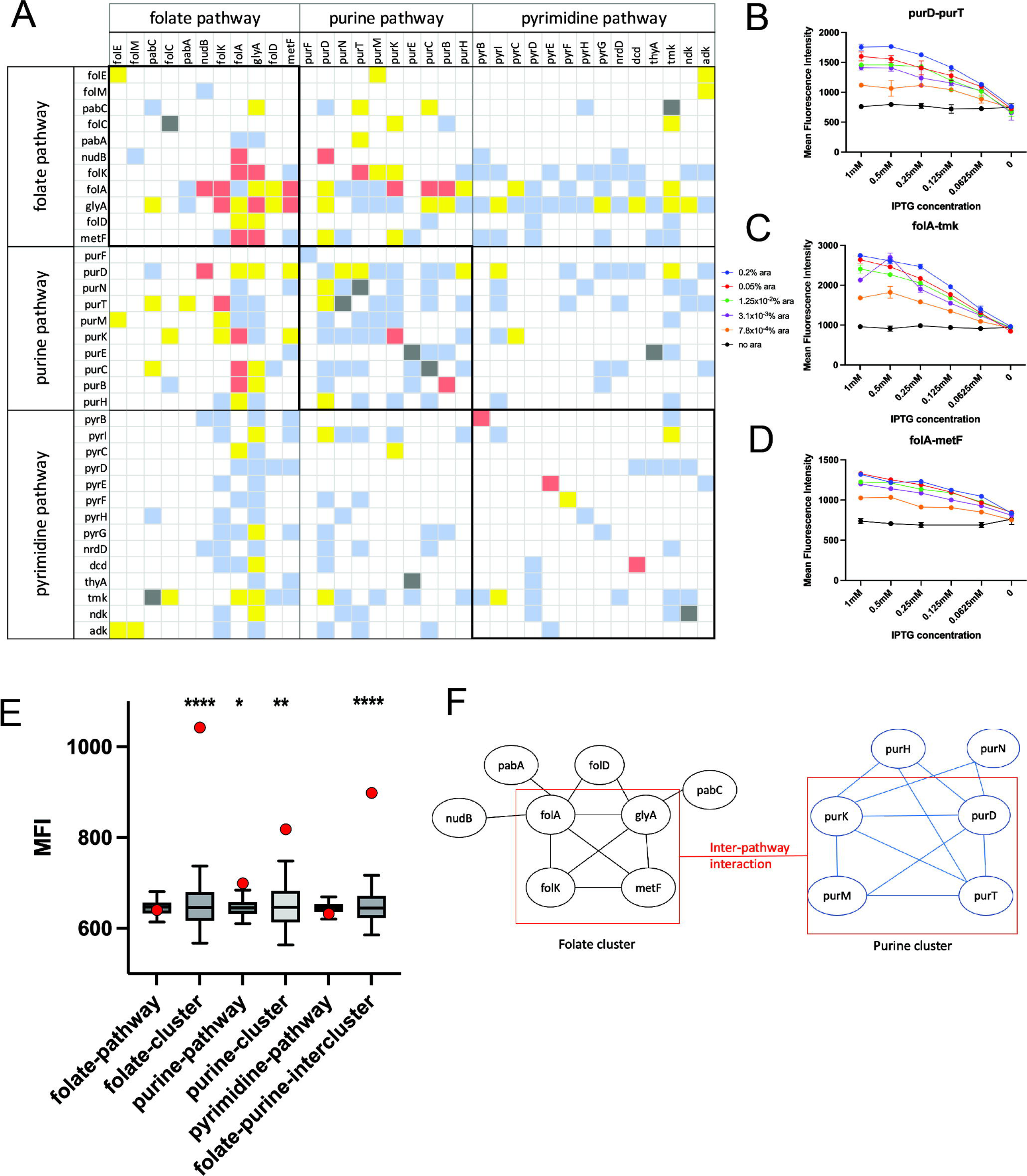
Exploration of the complete PPI network shows strong intra- and inter-pathway cluster of interactions. (A) A total of 35 proteins from the three different pathways were cloned as both N-terminal and C-terminal YFP fusions and interaction of every protein with every other protein (including itself which indicates homodimer probability) was probed using the split-YFP system. The matrix was color coded based on the mean YFP fluorescence of the pair. The interaction was evaluated in both formats (NYFP-A/CYFP-B and NYFP-B/CYFP-A) and the maximum fluorescence data for each pair was used to generate this matrix. Red boxes indicate MFI > 1100, yellow for 900 < MFI < 1100, blue for 750 < MFI < 900 while white boxes indicate MFI < 750. Titration of inducer levels (arabinose and IPTG) to test dependence of fluorescence values on intracellular protein expression for (B) purD-purT (C) folA-tmk and (D) folA-metF pairs. As expected, higher inducer concentration leads to more complex formation and stronger YFP fluorescence. Data is presented as mean of n=2 independent biological replicates, and the error bars indicate standard deviation (s.d.) (E) While there is no significant clustering for all 11 proteins in the folate pathway (n=55 pairs of interactions), a set of 5 proteins (n=10 PPI pairs for FolK, FolA, GlyA, FolD and MetF) show strong clustering. Similarly, a set of 5 proteins in the purine biosynthesis pathway (PurK, PurD, PurM, PurD, PurN) show cluster formation (n=9 PPI pairs). While no statistically significant clustering is observed among pyrimidine biosynthesis pathway proteins, there is strong evidence of inter-pathway interaction among selected folate and purine biosynthesis enzymes. For each cluster, the red data point represents the median MFI value of the cluster, while the box represents a null distribution of the median values of an equal number of interaction pairs (as the cluster size) randomly picked 10,000 times from the matrix. The box represents the middle 50% of the data and line in between represents the median of the distribution. The whiskers represent the 5-95 percentile interval. Statistical significance and p-value are calculated as described in Methods, *** indicates p-value < 0.001, ** indicates p-value < 0.01. No data normality is assumed. (F) A pictorial graph representation of the strongest interactions. Corners represent protein identity while edges represent strong interaction.

The BiFc method can yield false positives and the YFP truncated at positions 155/156 used in this study has been shown to have moderate tendency for self-assembly (11), hence we sought to benchmark the method in our case by comparing our results with two existing datasets.

First, surface plasmon resonance studies in our previous work (9) had showed that *E. coli* DHFR binds to PurH and GlyA, however shows no detectable binding to Adenylate Kinase (Adk). The split YFP data in this study shows high fluorescence for FolA-GlyA pair (1002) and for FolA-PurH pair (917), however much lower fluorescence for FolA-Adk pair (641), thereby corroborating the data obtained with purified proteins. Second, Fig 2A shows several high fluorescence values along the diagonal of the matrix. To find out if this accurately represents propensity of proteins for homo-oligomer formation, we plotted the fluorescence intensities for known cases of monomeric and oligomeric proteins (dimers and above). As shown in Fig EV2B, the distributions of MFI for known monomers and homo-oligomers were significantly different (p-value = 0.03; the differences become more strongly significant with p-value = 0.005 if the case of homo-dimeric PurT with MFI 175 is excluded from the distribution; given that in most cases, absence of interaction or weak interactions are characterized by MFI in the range of 500- 650, MFI of 175 seems more likely to be an outlier). Overall, Fig EV2B provides support that fluorescence complementation method in our study does not largely lead to false positives.

Since the N- and C-YFP fusion proteins are induced using IPTG and Arabinose, we next attempted to find out how the fluorescence intensity changes as a function of protein expression levels (or inducer concentration). As expected, we found that the increase in inducer concentration causes a sharp increase in YFP fluorescence (Fig 2B, C and D), indicating that the observed fluorescence are strongly concentration dependent.

### Folate and purine biosynthesis pathways show multiple strong intra- and inter-pathway interactions

We observed that certain proteins (FolK, FolA, GlyA and PurD) have a statistically higher propensity to be involved in PPI than others (Fig EV2D). Next, we sought to find out if folate, purine and pyrimidine biosynthesis pathways have significant intra-pathway interactions. Our results show that while the entire pathways do not have significantly stronger interactions than random pairs (folate, purine and pyrimidine pathways in Fig 2E), smaller clusters within the pathways do exhibit strong interactions that are statistically significant (folate cluster and purine cluster in Fig 2E). For example, five proteins of the folate pathway, namely FolK, FolA, GlyA, FolD and MetF interact strongly with each other, presumably forming an intra-pathway cluster (folate-cluster in Fig 2E, which refers to 10 PPI pairs among the five proteins listed above).

Interestingly, these are also sequential proteins in the folate pathway. Similarly, several proteins of the purine biosynthesis pathway (namely PurD, PurN, PurT, PurM and PurK) also form an intra-pathway cluster (purine-cluster in Fig 2E), though the interactions are overall weaker than those of the folate pathway. No such cluster was formed among enzymes from the pyrimidine biosynthesis pathway. The only exception in the pyrimidine pathway is Thymidylate Kinase (Tmk) which does interact with several proteins within its own pathway along with proteins from purine and folate pathways (Fig EV2D). Interestingly it was shown that Tmk overexpression is toxic, probably due to mis-interactions upon overexpression with other proteins of the folate pathway (9, 10). Next, we checked for significant inter-pathway interactions among the three pathways. While overall there are no statistically significant inter-pathway interactions when pathways are taken as a whole (Fig EV2C), certain clusters of proteins in the folate and purine biosynthesis pathway do show statistically significant inter-pathway interactions (16 PPI pairs referred to as folate-purine inter-cluster in Fig 2E). Based on the strong interactions observed, we constructed a graphical representation of possible clusters within the pathways and between folate and purine pathways (Fig 2F). According to this, FolA, FolK, GlyA and MetF form a core cluster in the folate pathway, where each protein interacts with every other protein. FolD, PabA, PabC and NudB proteins interact with one or two of the core proteins, but do not interact among themselves, hence they form peripheral members of the cluster. A similar analysis shows a core cluster in the purine biosynthesis pathway, comprising PurK, PurD, PurM and PurT, while PurH and PurN are peripheral members. We also calculated the statistical significance of the clusters identified in Fig 2E based on the original matrix of interactions in both configurations (Fig EV1A). As seen in Fig EV1B, the clusters still remained highly statistically significant, though the mean MFI of the cluster is lower. This is expected since the interaction may not be optimal in one configuration.

One possibility is that the pronounced number of interactions for certain proteins (eg, FolA, GlyA, FolK, PurD) observed in this work are due to other properties that are unrelated to functional interactions. For example, it might be that these subsets of proteins accumulate in inclusion bodies at the poles or simply have higher abundance than the other proteins, as YFP fluorescence is indeed strongly dependent on protein expression levels. To that end, we carried out a western blot analysis to quantify expression levels of the 35 N- and C-YFP fusion proteins (Fig EV3). The overall expressions of the CYFP-fusion proteins were less than NYFP fusions.

Though there were some variations in the expression levels within one set, they were not substantial enough to explain the variations in fluorescence intensities. For example, NYFP-FolA which shows strong interactions with multiple proteins (Fig 2A) had comparable expression to NYFP-PyrE and NYFP-Dcd, however PyrE and Dcd does not show any significant protein-protein interactions. On the other hand, NYFP-Ndk which has the highest abundance in the dataset shows poor interaction with most partner proteins. MetF which has low abundance in both NYFP and CYFP fusions, in fact shows strong interactions with several proteins in the pathway. Overall, this suggests that the high interaction propensity of certain proteins in the pathway is not due to their higher abundance.

Previous studies had clearly shown that over-expressed DHFR protein (FolA) does not accumulate in inclusion bodies to any substantial degree (12). To confirm this in the present study, we looked at the distribution of fluorescence intensity in live cell imaging experiments. Fig EV4 shows that in two representative strong interaction pairs (FolA-GlyA and PurD-PurT), the fluorescence was uniformly distributed across the cells with no accumulation at the poles, indicating that the observed YFP fluorescence is not biased due to protein localization at the inclusion bodies.

### Purine biosynthesis metabolon interactome is made up of structurally similar proteins

We next asked, are there physico-chemical characteristics of the enzymes that dictate their interactions? Theoretical studies have shown that structurally similar proteins tend to interact (13). To that end, we computed the structural similarity index of all proteins against all others using a structural similarity measure TM-score (14) (Fig EV5 and Table EV3). We first asked if the protein clusters identified from fluorescence data are structurally similar or not. We found that neither the entire folate pathway nor the 5 proteins in the folate cluster (FolK, FolA, GlyA, FolD and MetF, total 10 pairs) have a significantly higher median TM score compared to a null distribution (Fig 3A). On the contrary, the set of 5 proteins from the purine cluster (PurD, PurN, PurT, PurM and PurK, total 9 pairs) have a highly statistically significant greater TM-score distribution compared to the null distribution (p-value < 0.001) (Fig 3A), though the 10 proteins in the purine pathway as a whole does not share any significant structural similarity. The folate- purine inter-pathway cluster (16 interactions) also has no significant structural similarity. It is worth mentioning at this point that previous studies demonstrated that certain enzymes of the purine biosynthesis pathway evolved from a single common ancestor by gene duplication and divergence (15, 16). Thus, structural similarity within this subset of purine biosynthesis pathway proteins is likely to be a factor in selecting them as interacting cluster in the metabolon rather than a mere consequence of their origin as divergently evolved from a common ancestor.

**Figure 3:**
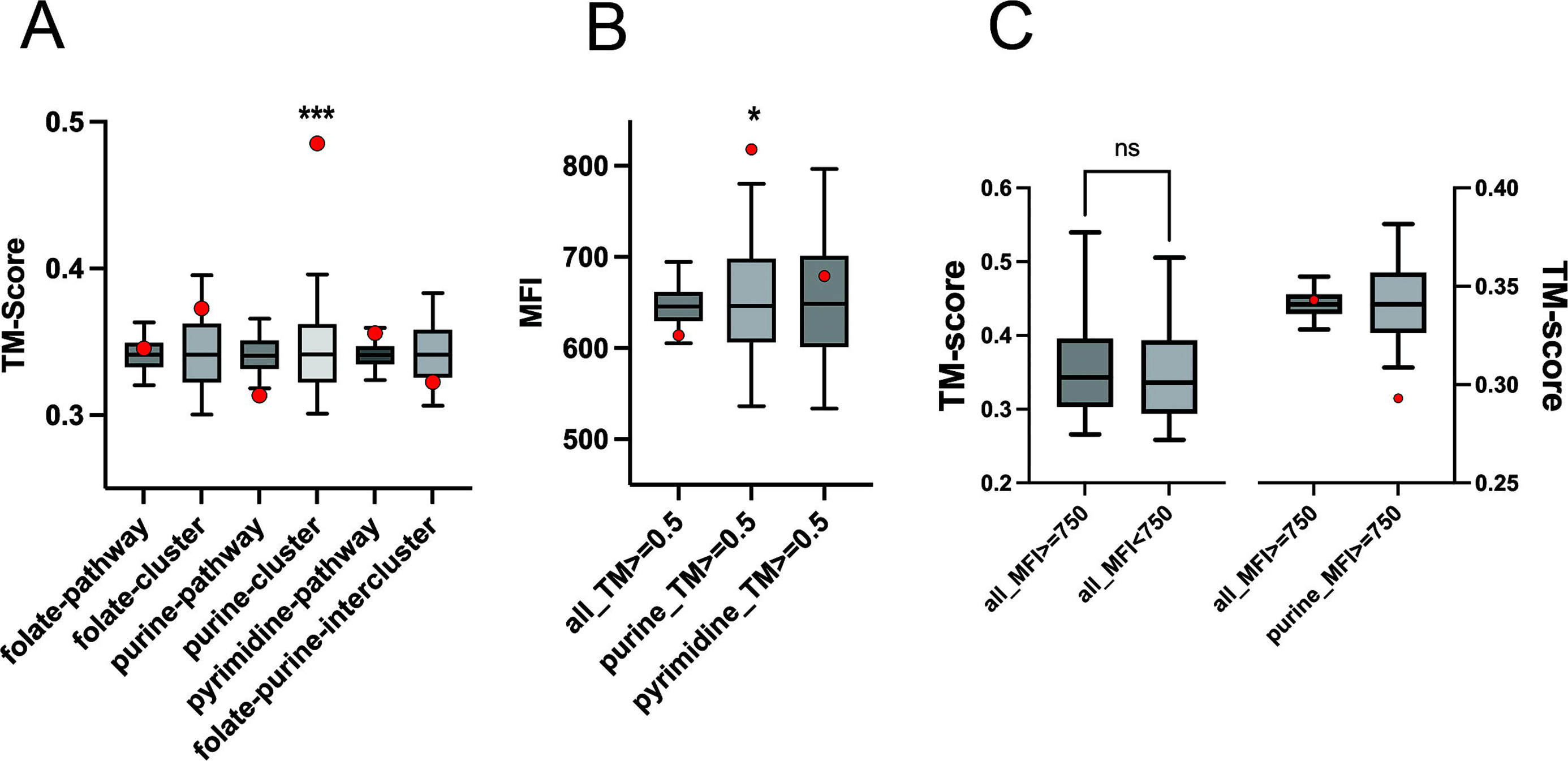
Subgroup of purine biosynthesis pathway proteins that interact strongly also have strong structural similarity. (A) shows median TM scores (red data point) for all groups presented in Fig 2E (all proteins in folate, purine and pyrimidine pathways, as well as the clusters) compared to a null distribution of TM values from the matrix (Table EV3). Only the purine cluster (PurD, PurN, PurT, PurM and PurK, total 9 pairs) have a highly statistically significant TM-score (*** indicates p-value < 0.001), while all none of the other pathway proteins show any significant structural similarity. The null distribution is generated from the median values of an equal number of interaction pairs (as the cluster size) randomly picked 10,000 times from the matrix (represented by the box plot, where the box represents the middle 50% of the data and line in between represents the median of the distribution. The whiskers represent the 5-95 percentile interval). Statistical significance and p-value are calculated as described in Methods. No data normality is assumed. (B) Median fluorescence intensities for all PPI pairs which show strong structural similarity (TM score > 0.5). The data shows that while high TM score by itself does not guarantee interaction, those in the purine biosynthesis pathway show mildly significant interaction propensity. The null distribution and p-values are calculated in the same way as panel (A). (C) A similar analysis showing median TM scores for all protein pairs that show strong interaction in the split YFP assay. The results indicate that structural similarity is not a general mechanism that leads to stronger interaction. On the left panel, the distributions of all_MFI>=750 and all_MFI=<750 are compared using a Student’s t-test, assuming normal distribution of data. On the right panel, like panel A and B, the red data point represents the cluster median, and the box plot represents the null distribution. p-values are calculated as in Methods.

To understand this further, we analyzed the fluorescence intensity of all those PPI pairs which have significant structural similarity (TM score > 0.5), both in the extended pathway as well as within individual pathways. Additionally, we also analyzed TM score distributions of all PPI pairs which have MFI values > 750 (Fig 3B, C). The results show that while a higher TM score overall does not guarantee interaction, it does dictate interaction propensity in the purine biosynthesis pathway.

### Saturation mutagenesis reveals dedicated PPI interfaces on DHFR that are conserved between its PPI partners

Since enzymes in a pathway interact via weak and transient interactions, no structural information is available yet for such PPI pairs by traditional methods. We reasoned that using our split YFP based methodology to screen for binding, we could highlight mutational hotspots in the PPI pairs that render these complexes stronger or weaker. To that end, we focused on two PPI pairs, FolA-MetF and FolA-FolK, which form one of the strong complexes in our dataset. We selected all surface exposed residues of FolA (accessibility > 80%, shown by pink spheres in Fig 4A), except its active site and contacting residues (within 5Å of the active site), and generated a near saturation mutagenesis library using VDS codon. VDS codes for all different amino acid types (hydrophobic, polar, charged), however does not incorporate large hydrophobics (F, W or Y) or proline, and does not incorporate any stop codon (Fig 4C). This is to ensure that the change in fluorescence (if any) will not arise due to protein destabilization or due to a truncated protein. When *E. coli* cells were co-transformed with NYFP-FolAlib and CYFP-MetF/CYFP-FolK (Fig 3B), the overall histogram of fluorescence intensity of the library variants did not appear to be different from WT (Fig EV6). On the other hand, single cells sorted from the top and bottom 5% of the distribution were found to have a large variation in fluorescence intensities. We selected 33 clones for the DHFR-MetF interaction and 28 clones for the DHFR-FolK interaction that had considerably higher or lower fluorescence intensities than the WT pair (Fig 4D, 4E), and sequenced them to reveal the identities of the mutants. Surprisingly, we found that a vast majority of the residues overlap between FolA surfaces interacting with FolK and MetF (26, 29, 114, 124, 131, 132, 140–145, 149, 157, 159) and are highlighted in Fig 4F and 4G. When these mutational hotspots were mapped back on to the structure of DHFR, we found the binding site is mostly located in the C-terminal beta hairpin, as well as in the helix close to the beta hairpin. The binding site does not overlap with the active site (folate or NADPH binding site).

**Figure 4:**
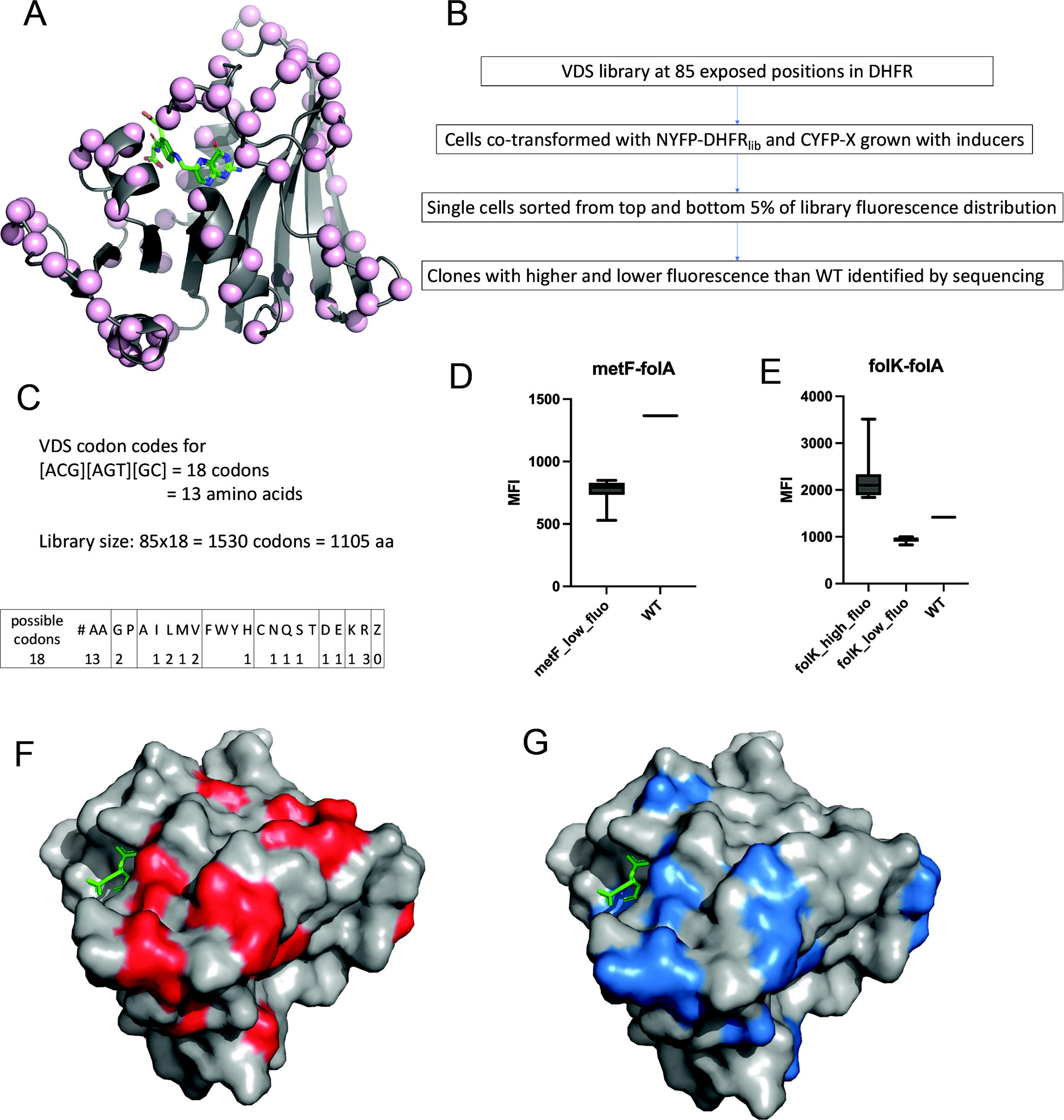
Elucidation of PPI binding interface using saturation mutagenesis-based library screening. At each of the 85 solvent exposed locations (% accessibility > 20%) in DHFR protein (Cα residue shown by light pink sphere in (A)), a VDS codon was introduced (one position at a time) by PCR. (C) VDS codes for 18 codons (13 different amino acids, no stop codons) which results in a total library size of 1105 amino acids. Panel (B) shows a flow-chart of the method used to map the binding interface. The NYFPlib plasmid was transformed with either CYFP-MetF or CYFP-folK expressing plasmids. Single cells were sorted out from the top and bottom 5% of the fluorescence histogram of the library variants, and clones with MFI significantly lower or higher than WT DHFR were sequenced to reveal their identities. Panels (D) and (E) represent boxplots of MFI for the sequenced clones. The box represents the middle 50% of the data and line in between represents the median. The whiskers extend from the minimum to the maximum values of the dataset. The mutations were mapped back on the DHFR structure and surprisingly, the two interaction surfaces show very significant non-random overlap (p-value=2e-7, see Methods and text) that is away from the folate binding site (DHF shown in green sticks). Panels F and G are DHFR structures where the MetF and FolK binding residues identified from the library screening are shown in red and blue colors respectively.

To test whether there is a non-random, statistically significant overlap between the FolA interaction surfaces with Folk and MetF (Fig 4F and 4G), we applied statistical test described in *Methods* and found that DHFR uses the same surfaces to interact with MetF and FolK: the overlap between two interaction surfaces is highly significant with a p-value of 10^-7^ (see *Methods*).

A possible alternative explanation for the drop in fluorescence in mutants is that mutations could lead to loss of DHFR protein abundance to unfolding/ misfolding/aggregation caused by DHFR destabilization. To address this possibility, we performed a western blot analysis of the mutants with anti-DHFR antibodies. None of the mutations showed any substantial change in intensity/abundance (Fig EV7), indicating that the mutations genuinely perturb binding without affecting abundance.

### Raman spectroscopy of purified proteins confirms binding site residues

Next, we sought to validate the interaction between DHFR and FolK (encoded by the gene folK) and its affinity changing mutants revealed in high throughput mutagenesis. To that end we performed Raman spectroscopy and specifically looked at the spectral features of the amide I range (1600 to 1700 cm^-1^) for DHFR and FolK individually and then upon mixing. Amide I spectra originate from the C=O stretch in the peptide backbone and are extremely informative about protein conformations. Analysis of Amide I spectra helps in the mechanistic understanding of secondary structures and their organization in a protein. Deconvolution of amide I spectra with standard peaks for discrete secondary structural elements viz. helices, sheets, and loops help in the quantification of secondary structure content in a protein (17). We performed deconvolution of the amide I spectra of the DHFR and FolK using Lorentzian peak fitting with pre-assigned peaks to better understand the change in the spectral features upon mixing both the protein solutions to detect non-additivity which would report on interaction between two proteins. The quality of fitting was checked with reduced chi-squared values, which were typically around 1.2.

Secondary structure contents obtained from deconvoluted DHFR (Fig EV8A) and FolK spectra (Fig EV8B) were compared with secondary structure contents of the crystal structures of DHFR (PDB: 1DRA) and FolK (PDB: 4M5I) to check the spectral quality and physical condition of the samples. Secondary structure contents for both DHFR (Fig EV8A) and FolK (Fig EV8B) were found to be comparable to the secondary structure contents of the reference structures.

Upon mixing the DHFR and FolK protein solutions any change in amide I spectral feature is a potential indication of spectra originating from the conformation of bound protein-protein complex which in turn indicates protein-protein interaction. In the absence of interaction, spectra appear as a mathematical sum of two individual protein spectra. In our study, we therefore investigated spectral features under conditions of DHFR FolK mixing. Control experiments were performed with ADK mixed individually with DHFR and FolK. We performed a similar quality check experiment with ADK as was performed with DHFR and FolK using the crystal structure of ADK as the reference and observed comparable secondary structure contents.

Upon mixing DHFR and FolK resultant amide I spectra showed new features around 1663cm^-1^ not observed in the individual spectrum of DHFR and FolK before mixing (Fig 5A). Further the resultant spectrum was not the mathematical sum of individual spectra obtained with DHFR and FolK before interaction (Fig EV9A). Interestingly, no new feature in the amide I spectra was observed for control acquisitions where ADK was mixed individually with DHFR (Fig EV9B) and FolK (Fig EV9C). The spectra obtained in these control acquisitions were equal to the mathematical sum of DHFR and ADK spectra before mixing in the DHFR-ADK mix and similarly FolK and ADK spectra in the FolK-ADK mix. The presence of new spectral feature with a peak around 1663cm^-1^ in the Amide I spectral profile of the DHFR-FolK mix (Fig 5A) is a potential indication of an interaction between FolK and DHFR and resultant Amide I spectra potentially reflecting the secondary structural organization of the FolK-DHFR complex in solution.

**Figure 5:**
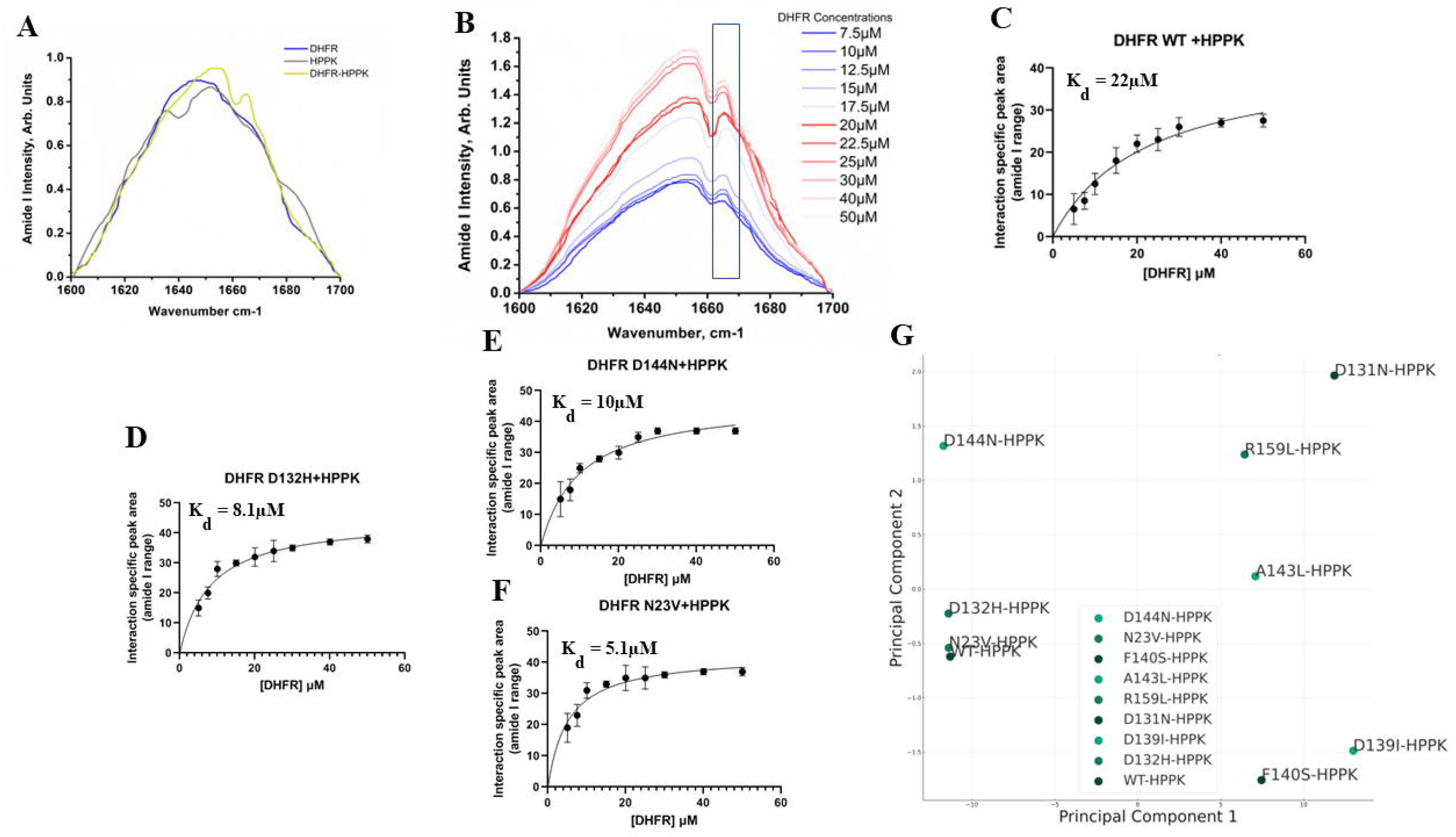
Raman Spectroscopic Analysis shows FolK interacts with WT and select mutants of DHFR. (A) Amide I Raman Spectroscopic profiles of WT-DHFR (blue trace), FolK (grey trace), and DHFR-FolK complex (yellow trace) are shown. Intensities (arbitrary unit, Arb. Units) for WT DHFR and FolK in the amide I range (1600-1700 cm^-1^) were recorded individually and then for the WT DHFR-FolK complex. The spectrum for the WT DHFR-FolK complex is not a mathematical sum of individual spectra recorded for WT DHFR and FolK and shows new spectral features and a characteristic peak at ∼1665 cm^-1^ indicating a signal originating from the WT DHFR-FolK bound complex. (B) Amide I spectral profiles of WT DHFR-FolK complex along an increasing concentration of DHFR (7.5 to 50 µM). The marked section shows the peak at ∼1665 cm^-1^ characteristic of WT DHFR-FolK interaction. (C) Changes in the peak area at ∼1665cm^-1^, characteristic of WT DHFR-FolK interaction, were plotted against increasing concentrations (7.5µM to 50µM) of DHFR. The peak areas were obtained upon deconvolution of the spectral read-outs (n=3) and the plotted data was subjected to Hyperbolic fitting to derive WT DHFR-FolK dissociation constant, K_d_ for WT DHFR-FolK interaction was found to be 22µM. (D) Changes in the peak area at ∼1665cm^-1^, characteristic of D132H mutant of DHFR-FolK interaction, were plotted against increasing concentrations (7.5µM to 50µM) of D132H DHFR. K_d_ for D132H DHFR-FolK interaction was found to be 8.1µM. (E) Changes in the peak area at ∼1665cm^-1^, characteristic of D144N mutant of DHFR-FolK interaction, were plotted against increasing concentrations (7.5µM to 50µM) of D144N DHFR. K_d_ for D144N DHFR-FolK interaction was found to be 10µM. (F) Changes in the peak area at ∼1665cm^-1^, characteristic of N23V mutant of DHFR-FolK interaction, were plotted against increasing concentrations (7.5µM to 50µM) of N23V DHFR. K_d_ for N23V DHFR-FolK interaction was found to be 5.1µM. (G) Principal component analysis (PCA) based on the entire Raman spectroscopic profiles (800 to 2000 cm^-1^) for WT and 8 mutants of DHFR. The positions of the mutants in the plot indicate their relatedness based on their Raman profiles. Mutants that are closely positioned on the plot share similarities in their Raman spectra. PCA shows FolK-interacting mutants, viz. D132H, N23V, and D144N along with the WT are clustered closely. On the other hand, non-FolK mutants of DHFR are separately spaced. n=3 biologically independent samples were used for the experiments. All the data are presented as mean values +/- SEM.

Next, we carried out Raman spectroscopy using a range of DHFR concentrations (from 7.5 to 50 µM) and keeping FolK concentrations fixed at 15 µM. We observed area under the characteristic peak at 1663 cm^-1^ increases monotonically (Fig 5B). The changes in the interaction-specific peak area at ∼1663 cm^-1^ were plotted against the increasing DHFR concentrations (Fig 5C). As the peak chosen was found to be specific for the interaction and no non-specific/unbounded fraction-specific signal was included in the plot, the hyperbolic model was used to fit the data. K_d_ was obtained upon hyperbolic fitting of the data and was found to be 22 µM for WT DHFR-FolK interaction (Fig 5C). Quality of the model deployed for fitting the data was assessed by R^2^ value and it was found to be 0.9. Next, we carried out similar interaction analyses with FolK for select mutants of DHFR that are predicted to weaken or strengthen binding based on FACS data (D144N, N23V and D132H increase fluorescence, while D131N, E139I, F140S, A143L, R159L showed loss in fluorescence). Interestingly, D132H mutant of DHFR did show stronger interaction with FolK with a K_D_ of 8.1µM (Fig 5D). Similarly compared to WT DHFR, D144N and N23V mutants of DHFR were also found to have a stronger interaction with FolK with K_D_ values of 10 and 5.1 µM respectively (Figs. 5E and F). For mutants that show a loss in fluorescence in the FACS analysis (D131N, E139I, F140S, A143L, R159L), quite strikingly, the characteristic peak at 1663 cm^-1^ was absent upon interaction, and hence an exact K_D_ could not be derived. While this directly hints at loss in binding, we attempted to compare more global signatures of the Raman spectra among mutants, hence we resorted to Principal Component Analysis (PCA). We performed PCA on the full Raman spectral profiles recorded across the 800–1900 cm ¹ range. This comprehensive spectral window includes not only the amide I region, which is commonly associated with secondary structural elements, but also broader backbone and side-chain vibrational features that can capture higher-order structural organization. PCA enables the reduction of this high-dimensional dataset into orthogonal principal components that account for the greatest variance, thereby facilitating an unbiased, global comparison of spectral signatures associated with each DHFR–FolK mixture.

In the resulting PCA projection (Fig 5G), samples corresponding to the wild-type DHFR–FolK complex and those involving high-affinity DHFR variants (D132H, D144N, N23V) form a coherent and tightly clustered group in PC1–PC2 space, indicating reproducible and structurally similar conformational changes associated with complex formation. In contrast, the profiles corresponding to DHFR variants previously shown to lose binding capacity (D131N, E139I, F140S, A143L, R159L) are dispersed away from this cluster, highlighting distinct spectral trajectories and supporting the absence of complex-specific conformational rearrangement. Importantly, this analysis does not rely on any single spectral feature—such as the emergence of the 1663 cm ¹ DHFR-FolK interaction specific peak—but captures cumulative differences across the entire spectrum, offering a robust validation of DHFR-FolK complex formation. PCA results not only confirm the presence of an interaction-specific signature in the DHFR–FolK complex but also establish that the resultant spectral profiles are not a mathematical sum of the components. PCA results provide an independent and comprehensive demonstration that the observed spectral patterns in interacting DHFR–FolK mixtures represent non-additive, interaction-specific conformational signatures, beyond what can be explained by simple peak-based analysis.

### High-throughput computational approach reveals that enzymes use similar interfaces to interact with multiple proteins

Mutagenesis-based library screening experiments so far reveal that the FolA protein interacts with two distinct partners, FolK and MetF with remarkably similar interaction interfaces. However, due to experimental limitations, the interaction interface study could only be performed for a limited number of interaction partners. To study whether multiple folate pathway proteins use similar interface to interact with each of their PPI partners, we employed a high-throughput computational approach using AlphaFold3. We first use AlphaFold3 to predict dimeric structures, for all pairwise interactions between different members of the folate pathway (Fig 6A), like the experimentally generated matrix in Fig 2A. Subsequently, we performed metadynamics simulations (see detailed description in Methods) to further refine these dimeric structures and their corresponding interaction interfaces (Fig 6A). To elucidate the most stable PPIs, we identify pairwise inter-chain residue-level contacts for all structures that correspond to free energy minima identified in the metadynamics simulations (Fig 6A). We also compute binding free energies (ΔG_binding_) from the free energy profiles obtained using our metadynamics simulations. In Fig 6B, we compare whether the protein pairs computationally predicted to be stable binders (ΔG_binding_ < 3 kcal/mol) also show stable binding in experiments. For the sake of comparison, dimers predicted to be stable binders in simulation were considered as “true predictions” if the MFI for the corresponding protein-pair in the fluorescence experiments were greater than 670. We note that more than 65% of the simulation predictions are “true predictions” for a range of ΔG_binding_ and experimental fluorescence thresholds (also see Fig EV10). The robustness of simulation predictions is consistent across the three pathways, and for a range of experimental and simulation thresholds (Fig 6B and Fig EV10). These results suggest that our metadynamics simulation protocol can be reliably employed to screen PPIs and identify stable binders. It must be noted that such an approach for comparing experimental fluorescence values was chosen over a direct correlation analysis between the binding free energies from simulation and fluorescence because the two techniques could have very different dynamic ranges. In other words, an increase in the ΔG_binding_ computed from the metadynamics simulations might not directly correlate to a commensurate increase in fluorescence values in experiments making the exercise of computation of a direct correlation somewhat misleading. The simulation data is for simple two component binding, unlike the experimental values which could be perturbed by presence of other molecules in solution. Therefore weak, stable binders in metadynamics simulations might not directly show up in fluorescence signals in experiments at biologically relevant concentrations in a more complex cellular milieu. The free-energy minimum structures from our simulations are further used for analysis of interaction-interfaces.

**Figure 6:**
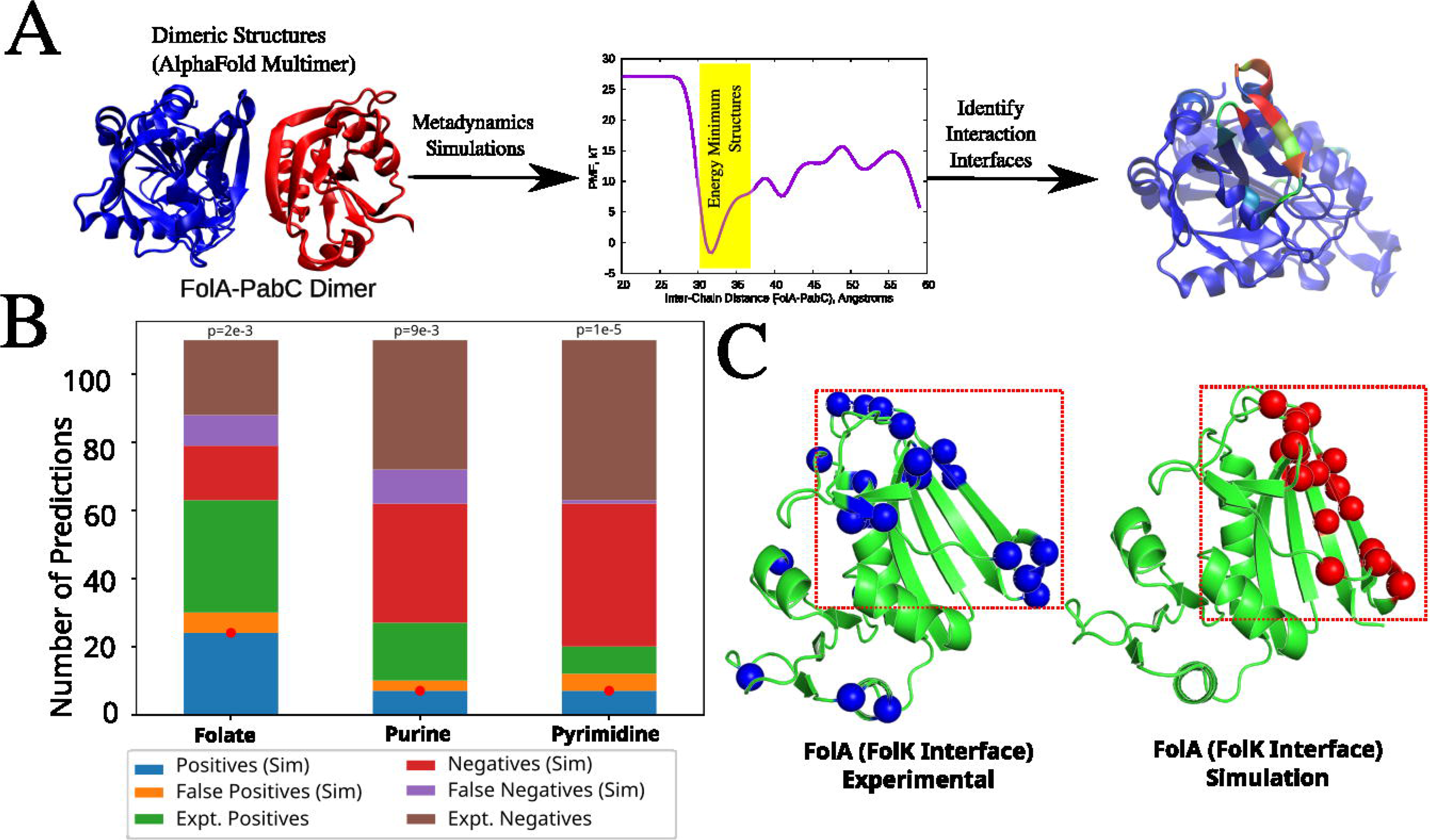
Metadynamics simulations used to compute interaction interfaces and unbinding free energies for folate pathway dimers. A) AlphaFold 3.0 is used to make initial dimeric structure predictions for all possible folate pathway enzyme pairs. To further refine these structures and compute their stabilities, metadynamics simulations were performed, with AlphaFold3 predicted structures as starting points and with inter-chain distance as the collective variable. The structures corresponding to the free energy minima in the PMFs were then used to study the interaction interfaces for folate pathway proteins. B) Dimers with ΔG_binding_ < -3 kcal/mol in metadynamics were considered stable binders. Those dimers predicted to be stable binders in simulation were considered as “true predictions” if the MFI for the corresponding protein-pair in the fluorescence experiments were greater than 6**7**0. p-values above each bar indicate statistical significance of agreement between theory and experiment against null hypothesis that simulations predict binding surfaces between pairs of proteins at random. A more complete analysis of agreement between computational predictions and experiment for a range of thresholds is in Fig EV10. C) Simulations and experiments predict similar interaction surface for FolA with FolK. Residues of FolA involved in interactions with FolK based on experimental data (marked as red beads in left structure) and metadynamics simulations (marked as cyan beads in right structure). The simulation predicted surface and the experimentally detected interaction surfaces show significant overlap (p-value of 2.7x10^-5^ for the null hypothesis that experimental and predicted interaction surfaces have random overlap, see Methods for detail).

In Fig 6C, we then compare the extent of overlap in predicted interaction interfaces of the FolA (from metadynamics simulations) for the FolA-FolK pair and compare it to the experimentally determined interface. Fig 6C shows that the simulations predictions agree with the experimentally predicted interaction interfaces, suggesting that simulations can be used to screen interaction interfaces in a high-throughput fashion. The overlap between the experimental and simulation predicted surfaces is statistically significant with a p-value of *2.3x10^-5^*, under null hypothesis that computation and experiment pick two unrelated surfaces on FolA. (see Methods) Interestingly, structural alignment of the AlphaFold3 predicted complexes of FolA with its strongest partner proteins NudB, FolK, GlyA and FolD clearly shows that different proteins bind around a similar region of FolA (Extended View Fig EV11A). To gain a more quantitative insight into this observation for all folate pathway proteins, we calculate the frequency of involvement of every amino acid residue in heteromeric PPIs across different partners within the pathway. In Fig 7A, we represent the AlphaFold3 predicted structures of folate pathway proteins with residues that are frequently involved in PPI across different partners represented as large (yellow to green colored) spheres while those which are less involved in PPIs are represented as smaller, darker colored spheres. As seen from Fig 7A, the residues most frequently involved in hetero-dimeric protein-protein contacts across all their PPI partners are clustered in a small region of the protein, as evident from the spatial segregation of bright and dark colored spheres on the folate pathway proteins in Fig 7A.

**Figure 7:**
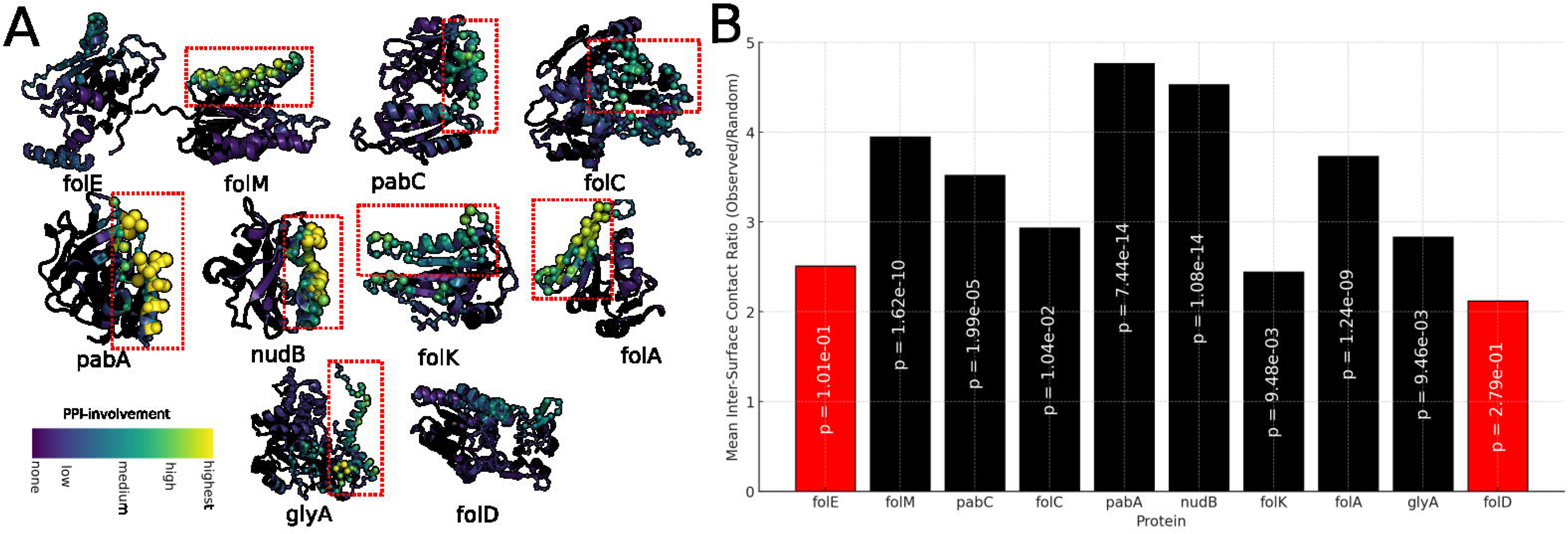
Folate pathway proteins use dedicated conserved surfaces to interact with all their metabolon partners. A) Representation of amino acid residues that are most frequently involved in hetero-dimeric protein-protein interactions in the folate pathway. 10 folate pathway enzymes are represented in the figure, and the most frequently observed residues in PPIs are represented by spheres. The radii and color of the spheres are represented such that most residues that appear most frequently in PPIs (in simulation) are represented by larger spheres that are brightly colored. Smaller, dark shaded spheres indicate residues that are not frequently involved in PPIs with other proteins. B) The ratio of observed inter-surface contacts to inter-surface contacts expected under the null hypothesis for randomly occurring surfaces. This ratio is presented for all 10 folate-pathway proteins. p-values vs random overlap null model (see Methods) is shown for each protein.

To check if there is a non-random overlap of interaction surfaces across different interaction partners, we employ the statistical analysis (see Methods section for details) which provides p-value of observed overlaps between interaction surfaces with different interaction partners against the null hypothesis that a protein utilizes unrelated surfaces to bind to different interaction partners in the folate metabolon (also see Extended View Fig EV12). In Fig 7B, we present the results of this analysis in the form of the ratio of mean inter-surface contacts for the observed surfaces (from metadynamics simulations) versus that of control pairs of randomly drawn surfaces on the protein. A high ratio in Fig 7B suggests that the observed surfaces share a high degree of overlap compared to what would be expected for randomly chosen surfaces on the protein. As evident from the p-values shown in Fig 7B, we observe a statistically significant overlap in interaction surfaces across different interactions for all proteins of the folate pathway, except FolE and FolD. The corresponding Z-scores are presented in Extended View FigS12. Interestingly, despite being involved in interactions across multiple different partners, similar interfaces on the protein get shared across several different folate-pathway interaction partners. The metadynamics simulations, therefore, suggest that these proteins could exhibit a great deal of promiscuity in PPIs, with the same interface being involved in interactions across multiple different partners. A remarkable exception seems to be FolD, which despite being involved in strong interactions with FolA and GlyA, does not seem to involve a common interaction interface. Structural alignment (Extended View Fig EV11B) shows that FolA and GlyA bind on two opposite surfaces of FolD.

Lastly, since several proteins in the folate as well as 1-carbon metabolism pathway are homo-oligomers (as evidenced by the high fluorescence values along the diagonal), we asked if the surfaces mediating weak PPIs have the potential to disrupt the oligomeric interface. To that end, we looked at the homo-dimeric as well as the hetero-dimeric structures of GlyA and PurK, which show strong homo-oligomer signals as well as PPI with multiple proteins. Interestingly, in a few representative cases that we analyzed (FolA:GlyA in folate pathway; PurD:PurK in purine pathway; and FolA:PurK in the folate-purine inter-pathway), the interface mediating weak PPI were completely different from the oligomeric interface (Extended View Fig EV13), indicating that these are truly novel interaction interfaces predicted by AlphaFold3 that do not rely on the sticky hydrophobic patches on the oligomeric interfaces.

### Folate-pathway metabolons show a dramatic increase in metabolic flux for the pathway compared to monomeric enzymes

Our experiments and simulations so far show the presence of protein-protein interactions between enzymes of the same pathway. To verify that interaction networks derived from fluorescence experiments result in formation of spatial enzyme clusters, we employ a Coarse-Grained (GC) model where the enzymes are modeled as patchy particles. Each enzyme in the Langevin dynamics simulations is modeled using a central hard sphere of radius 10Å and two patches on the surface of this hard-sphere corresponding to a PPI-site and a substrate-binding site (Fig 8A). The substrate-binding and PPI-site patches have 10 distinct identities to account for the 10 enzyme types of the folate pathway being modeled here (see Fig 8A where we show the different coarse grained patchy particle enzymes). 5 independent trajectories of 10 microseconds each were used to study the functional dynamics of this multi-enzyme system. As seen from Fig 8B, the enzymes form dense dynamic clusters that are stable at simulation timescales. Interestingly, even a relatively simple PPI energetics based on the broad MFI ranges discussed previously, results in a dense PPI map observed in the simulations (Fig 8C). The cluster size distribution at equilibrium (Fig 8D) shows an exponential-like dependence and presence of several stable large clusters. These results show that a non-isotropic interaction model with a single interaction surface shared with multiple partners and with protein-protein interaction energetics derived from the fluorescence experiments (Fig 2), can support the formation of transient enzyme clusters within the folate pathway.

**Fig 8.**
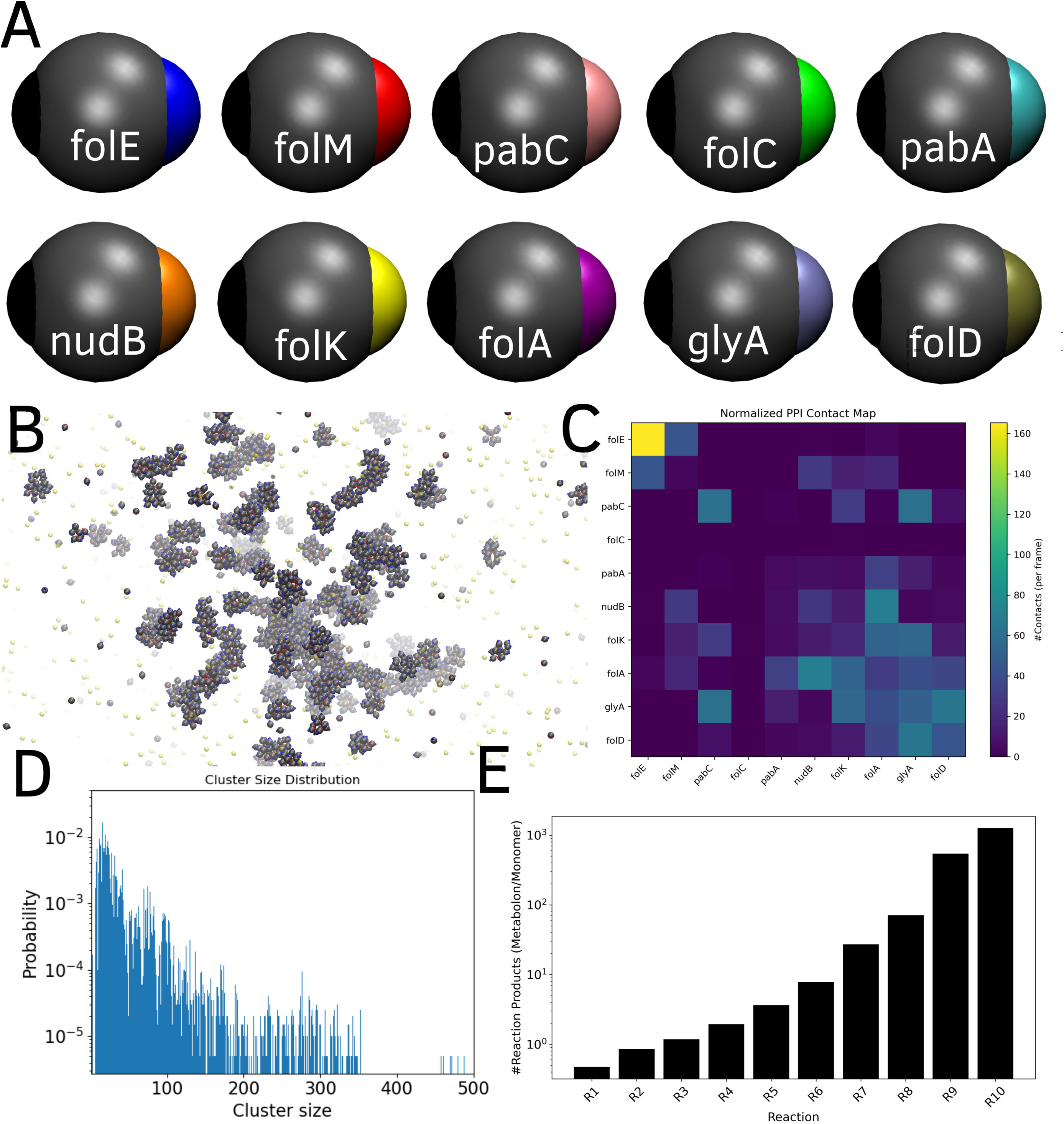
Reaction-diffusion Langevin dynamics simulations of the folate pathway. A) Patchy particle model describing the 10 folate pathway proteins. B) Formation of enzyme clusters during the 1 μs Langevin dynamics simulations. C) Mean numbers of protein-protein contacts between different coarse-grained enzyme particles in the Langevin dynamics simulations. D) Size distribution of enzyme clusters at equilibrium. E) Reaction fluxes for each reaction in the metabolon state normalized by the same quantity in the monomeric (unclustered) state of the enzymes in the control simulation where PPI are abrogated. The initial reactions in the pathway (R1 to R4) show either a slight drop or no change in flux upon clustering. Later reactions in the pathway, however, show a dramatic increase in fluxes approaching several orders of magnitude showing the net gain in efficiency of substrate channeling in longer pathways upon clustering of enzymes.

The results so far show that stable enzyme clusters could be supported for a heterogenous interaction map where only a subset of the enzyme pairs show stable interactions. How does the formation of dynamic clusters affect the net metabolic flux of the folate enzyme pathway? To address this question, we introduce a diffusion-reaction protocol in our LD simulations such that the 10 patchy particle enzymes in the simulation catalyze 10 sequential reactions in a linear reaction cascade (see (18) and Methods section for more detail).

In this study, we performed the diffusion-reaction simulations for 2 scenarios -

1. No protein-protein interactions are imposed ensuring the enzymes are in a monomeric state, and all reactions occur in the bulk.
2. Energetics of PPI between enzymes is derived from the experimental PPI-matrix (Fig 2) corresponding to the folate pathway (Fig 8C).

5 independent 10 microsecond trajectories were simulated for both scenarios. The output of each reaction is tracked at the end of the simulation. In Fig 8E we plot the ratio of the number of successful reactions in the clustered state for enzymes normalized by the same for monomeric enzymes. Interestingly, as we move along the pathway (R4 and higher in Fig 8E), the reaction fluxes show a significant gain for clustered enzymes, at the expense of a slight drop in flux for the initial reactions of the pathway. The gain in efficiency for later steps in the pathway can be several orders of magnitude compared with than for the monomeric unclustered state.

Our results, therefore, suggest that experimentally determined PPI between enzymes of the folate pathway results in enzyme clusters which can improve the efficiency of metabolic turnover of the pathway. The reaction probabilities chosen in this study are drawn from a realistic distribution of kcat/KM that peaks several orders of magnitude lower than the diffusion-limit Fig EV15). Therefore, our results strengthen the hypothesis that clustering of enzymes via PPIs can improve metabolic fluxes of pathways made up of otherwise “imperfect” enzymes [Fig EV16 and (18)] which operate far from the diffusion limit. Crucially, simulations with the patchy particle model show that even when the enzymes can interact via a single interface shared across several partners, the enzymes can exist in a clustered state that can support several orders of magnitude higher fluxes for the pathway compared to runs occurring in bulk solution. The stoichiometries of enzyme assemblies that form via the folate pathway PPI in our simulations (Fig EV17) reveal that all the enzymes of the pathway are enriched to the same extent in large clusters.

## Discussion

The one-carbon folate biosynthesis pathway comprising three intersecting and inter-dependent pathways (de novo purine, pyrimidine and methionine synthesis) is one of the central metabolic networks of the living cell. It has been long hypothesized that metabolon formation may be essential to this pathway to protect labile folate co-factors that are easily prone to oxidative damage (19). In the absence of reducing agents, folate derivatives were shown to have a half-life of a few minutes in vitro (20), however they appear much long-lived in *in vivo* studies (21, 22). Moreover, the fact that several mammalian proteins in the 1-carbon metabolism pathway had more than one binding sites for folate co-factors indicated that substrate channeling was potentially inherent to this pathway (23). In our earlier work, immunoprecipitation followed by mass spectrometry showed extensive albeit weak and transient protein-protein interactions between DHFR and its functional neighbors (9). In this work, we systematically delineate the extent of protein-protein interactions in the entire *E. coli* 1-carbon metabolism pathway, using a bimolecular fluorescence complementation assay that can capture transient interactions in vivo. We identified two clusters of proteins in the folate and purine biosynthesis pathways respectively that had very strong intra-cluster as well as inter-cluster interactions.

A remarkable finding of our study was that the extent of interaction was the strongest among members of the folate biosynthesis pathway followed by those in the purine biosynthesis pathway. Based on our data, there was no statistically significant interaction among the pyrimidine biosynthesis pathway proteins, though in several cases, they formed strong homomers (along the diagonal in Fig 2A). This trend in interactions aligns quite strikingly with the order of stabilities of the metabolites in the respective pathways as folate metabolites (5,10- methelene THF, 5-methyl THF, N-formyl THF) are considered the most labile (20). On the other hand, pyrimidine pathway intermediates are mostly stable metabolites in the form of mono- phosphates or tri-phosphates (OMP, UMP, UTP, CTP, dTMP, etc.) and presumably do not need metabolon like networks to protect them in the cytosol, while purine pathway intermediates are less stable (GAR, FGAR, AIR, NCAIR, SAICAR), as they are often difficult to quantify from the cell lysate using metabolomics (10, 24). This observation may also hint more generally towards the possible evolution of PPI networks and metabolons in enzymatic networks to protect labile metabolites. Spatial clustering of functionally related enzymes into a dense cluster could also result in significant increase in metabolic flux compared to the bulk reaction scenario, as previously demonstrated by mechanistic models (18) (25)

Though much evidence is present in the literature about the existence of metabolons in various organisms, very little information is available about structural architecture of these complexes (8, 26, 27), particularly the nature of these PPI interfaces and their binding affinities. Some interesting questions are: do multiple proteins in a metabolon associate together, or do binary interactions occur at a time? How do these weak interaction interfaces differ from obligate PPI? Are active sites of the enzymes predominantly involved in the interaction interface? A key highlight of the current study is that it provides structural insights into the structural detail of the 1-carbon metabolon using a combination of high-throughput mutagenesis and screening experiments followed by meta-dynamics simulations in combination with AlphaFold3 prediction of initial complexes. Not only did the calculations predict PPIs with high accuracy as well as predict interfaces that agree with experimental findings, but they also shed light on some remarkable and unanswered aspects of the extended folate pathway metabolon. First, it shows that proteins that interact with multiple partners in the pathway almost always use the same interface for interaction with multiple metabolon partners, indicating a high degree of promiscuity at the interaction interface. This also suggests that monomeric enzymes are presumably involved in only binary interactions at a time. However, for oligomers, multiple subunits can associate with different proteins, and hence higher order complexes may form, though we do not have such evidence so far from our present study. Second, our data on DHFR also shows that the PPI interface most used by the enzyme is far away from the active site. Proteins that interact with multiple partners (the so-called hub proteins) can be of two classes: date hubs (singlish hubs) wherein pairwise interactions happen one at a time, or party hubs (multi-hubs), which can engage multiple partners at the same time (28, 29). Our findings suggest that promiscuous hub proteins in the 1-carbon metabolism pathway behave more like date hubs. Whether this is a general phenomenon for pathway enzymes warrants further investigation. For example, in the only other study where some degree of structural information is available for metabolons (8), cross-linking followed by mass spectrometry showed that citrate synthase of TCA cycle uses two distinct PPI interfaces to interact with malate dehydrogenase and aconitase, akin to FolD in our study. As observed in this study, the PPI interface was distinct from the active site. In case of purinosomes (4), a luciferase reporter-based Tango reporter assay to assess PPI revealed that three enzymes hPPAT, hTrifGART, and hFGAMS of the purine biosynthesis pathway form a core structure, while other proteins associate on to them. In the absence of any structural information, it is unknown if the enzymes use a common interface or can associate simultaneously, hence future work will establish if our observations are a general property of metabolons.

Nevertheless, one might surmise as to why evolution might favor singlish hubs over multi-hubs, as date hubs ensure that one interaction must have to dissociate for another new interaction to happen, and therefore usage of the same PPI interface might be important for the dynamic assembly and disassembly of interactions that is so fundamental to the concept of metabolons. Shared interfaces might also pose a natural limit on the size of the metabolon, which otherwise can form giant assemblies that completely sequester proteins. It can also act as a density modulator that can increase metabolic fluxes in long biochemical pathways (18).

Our finding about the *E. coli* 1-carbon metabolon is one of the few examples of metabolons in prokaryotic cells. Much work in the recent past have shown that biochemical reactions do not take place in an unorganized manner in the cytosol of bacteria, rather there is a considerable degree of self-organization inside them. Other examples of such supramolecular enzyme clusters in bacteria include the TCA cycle pathway in *Bacillus subtilis* (30) and the glycolysis pathway in *E. coli* (31).

Our previous work on *E. coli* DHFR (9) suggested that one of the most important fitness consequences of metabolons is that when individual components are over-expressed, it sequesters others in the non-physiological complex forming near permanent interactions, thereby causing toxicity to the cells. A more detailed study on the propensity of a protein to be involved in PPI and its over-expression toxicity on a much larger dataset of proteins could shed light on sequestration of partner proteins through interaction as a general mechanism of toxicity in metabolic enzymes.

One might argue that our study merely shows interactions among multiple components of the pathway and did not present evidence of substrate channeling. Zhang and Fernie (7) argue that only enzyme assemblies that are shown to both physically and functionally interact (channel metabolites) should be termed metabolons. However, considering that we observe the strongest interaction among enzymes with the most labile and short-lived substrates (folate pathway), and no interaction among the pyrimidine pathway with the most stable intermediates, suggest that substrate channeling must have been an evolutionary driving force to form transient complexes to protect unstable ligands from degradation by limiting their diffusion paths in the cytosol. Furthermore, our dynamic modeling of the folate pathway with and without formation of metabolon showed that metabolon formed via experimentally observed PPI speeds up the flux through the pathway by several orders of magnitude due to substrate channeling. Therefore, the modeling results highlight clear biological significance of the observed PPI in the folate pathway. This speed up effect is especially pronounced for longer pathways such as folate pathway in this study as the advantage from metabolon formation is less pronounced (and actually could be reversed in some cases) for shorter pathways comprised of 2-3 reactions (18). Crucially, the increased fluxes for the pathway in our simulations was purely an outcome of protein-protein interactions without an explicit substrate channeling mechanism incorporated. PPIs could therefore have evolved to optimize biochemical yields for the whole pathway while individual enzymes could operate far from the diffusion limit. A more explicit substrate channeling mechanism could further enhance the efficiency of enzyme clusters beyond that achieved by a simple clustering of enzymes via PPIs. Whether enzyme clusters that are much smaller than phase-separated droplets or condensates can sufficiently enhance reaction fluxes is a natural question that arises considering the findings from our current study. The shape, size, stoichiometry and dynamic properties that result as an outcome of shared protein-protein interaction surfaces as opposed to multiple interaction patches would be critical to holistically understand the biological implications of the current work. Future work with more detailed insight into the effect of clustering on metabolic fluxes will decisively answer the outstanding questions concerning functional and evolutionary significance of metabolons.

## Acknowledgement

This work was supported by NIH grant R35GM139571 to E.I.S. The authors thank the flow cytometry facility (Bauer core) and the Harvard Center for Biological Imaging core at Harvard University.

## Author contributions

S.B and E.I.S conceived the study; S.B and S.R designed research; S.B, S.R, S.C, B.V.A, M.K performed research; S.B, S.R, S.C, B.V.A, E.I.S analyzed data; S.B, S.R, S.C, B.V.A, E.I.S wrote the paper.

## Declaration

The authors declare no competing interests. The work was entirely performed when authors were part of Harvard University.

## Extended View Figure legends

**Figure EV1:** (A) The complete asymmetric matrix of all interactions measured in both configurations (NYFP-A/CYFP-B) and (NYFP-B/CYFP-A). Fig 2A is derived from this matrix by taking the maximum value for each PPI pair. As in Fig 2A, red boxes indicate MFI > 1100, yellow for 900 < MFI < 1100, blue for 750 < MFI < 900 while white boxes indicate MFI < 750. Grey boxes indicate no measurement. (B) Based on the original matrix in panel A and Table EV2, we calculate the statistical significance of the clusters identified in Fig 2E. The results are largely similar to Fig 2E. For each cluster, the red data point represents the median MFI value of the cluster, while the box represents a null distribution generated from the median values of an equal number of interaction pairs (as the cluster size) randomly picked 10,000 times from the matrix. The box represents the middle 50% of the data and line in between represents the median of the distribution. The whiskers represent the 5-95 percentile interval. Statistical significance and p-value are calculated as described in Methods, *** indicates p-value < 0.001, ** indicates p-value < 0.01. No data normality is assumed.

**Figure EV2:** (A) Frequency distribution of mean YFP fluorescence intensities of 1225 interactions from the dataset. The intensities show a wide range of variation. (B) All proteins in our dataset for which oligomeric status is known from crystal structure data (total 30 proteins) were grouped into two classes – monomer and oligomer (dimer and above). The plot shows distribution of MFI values of all proteins belonging to that group. An unpaired student t-test shows that oligomers have significantly higher fluorescence intensity compared to monomers (p-value = 0.03, n=10 for monomer set, and n=20 for oligomer set). The red data point corresponds to PurT, for which the MFI of 175 was much lower than the cellular background fluorescence for non-interacting protein pairs. If this data point is excluded, then the p-value becomes strongly significant with value of 0.005. (C) There is no significant inter-pathway interaction among folate, purine and pyrimidine biosynthesis pathways when entire pathways are taken into consideration. For each cluster, the median MFI value of the cluster (red data point) is compared against a null distribution that is generated from the median values of an equal number of interaction pairs (as the cluster size) randomly picked 10,000 times from the matrix (represented by the box plot, where the box represents the middle 50% of the data and line in between represents the median of the distribution. The whiskers represent the 5-95 percentile interval). Statistical significance and p-value are calculated as described in Methods. No data normality is assumed. (D) Interaction propensity of each protein with all other proteins in the dataset represented as average of MFI across all 35 proteins in the dataset. The median (red data point) of all 34 interactions of a protein (except with itself) is compared against a null distribution, as explained in panel C.

**Figure EV3:** Western blot images to check expression of (A) NYFP- and (B) CYFP- fusion constructs using polyclonal anti-YFP antibody (see Methods for details). Since NYFP fragment is larger, NYFP-fusion proteins have higher intensities on the blot than their CYFP counterparts, and therefore blot (B) shows significantly more background. In all cases, the most prominent band that matches the expected molecular weight of the fusion proteins was used for quantification. Overall, though expression levels of the fusion proteins vary, they are not sufficient to explain differences in observed PPI strengths.

**Figure EV4**: Phase contrast images superimposed with YFP fluorescence of live *E. coli* cells that express (A) NYFP-FolA & CYFP-GlyA and (B) NYFP-PurD & CYFP-PurT. In both representative cases where we observe high fluorescence from FACS, the corresponding images show that YFP fluorescence is uniformly distributed inside the cells, as opposed to localization at certain areas which might signify protein aggregation.

**Figure EV5:** A 35x35 matrix for TM score of each protein pair. A symmetric matrix was generated by taking the maximum value for each protein pair.

**Figure EV6:** Fluorescence intensity distributions of ∼30,000 cells transformed with NYFP-WT DHFR/CYFP-MetF plasmids (left panel) and NYFP-VDS_lib DHFR/CYFP-MetF plasmids (right panel). The histogram obtained with VDS library does not appear to be different from the WT histogram.

**Figure EV7:** Protein expression check using western blot for DHFR library variants which show substantial loss or increase in fluorescence relative to WT. No significant change in expression was observed, indicating that the change in fluorescence is not due to generic loss of protein levels.

**Figure EV8: (A)** Amide I (1600 to 1700 cm^-1^) Raman spectral profile of WT DHFR. The plot shown is the mean of the acquisition from three independent technical repeats. The bar plot shows the secondary structure contents of DHFR (black bars) as obtained upon deconvolution of the Amide I spectrum. Amide I spectra for DHFR were deconvoluted using Lorentzian peak fitting and the secondary structural contents for helix, beta sheet, and turns and loops were compared with the solved structure (Black bars, PDB- 1DRA). Peaks were selected based on the standard Amide peaks used for secondary structure analysis and the quality of the fit was checked using reduced chi-squared value which was typically between 1.2 to 1.5. The secondary structural content was found to be comparable to the PDB structure, further reflecting the quality of the spectra and the peak fitting carried out. **(B)** A similar analysis was done for FolK. The secondary structural content was found to be comparable to the PDB structure. **(C)** Amide I spectral profile of the control protein ADK was acquired. A similar analysis was done for ADK, and the secondary structural content was found to be comparable to the PDB structure. n=3 biologically independent samples were used for the experiments. All the data are presented as mean values +/- SEM.

**Figure EV9:** (A) Raman Spectra obtained from DHFR-FolK interaction set do not overlap with the hypothetical spectrum, which is the mathematical sum of DHFR and FolK spectra. Raman Spectra for control sets using ADK and each of DHFR (B) and FolK (C) were acquired using laser excitation of 785nm. Intensities (arbitrary unit, Arb.Units.) for native DHFR and ADK in the amide I range (1600-1700 cm^-1^) were recorded individually and then upon mixing. The resultant spectra obtained were found to be comparable to the mathematical sum of the individual spectrum of DHFR and ADK, potentially suggesting the absence of any interaction. (C) The resultant spectrum was found to be comparable to the mathematical sum of the individual spectrum for ADK and FolK which suggests absence of any interaction. n=3 biologically independent samples were used for the experiments. All the data are presented as mean values +/- SEM.

**Fig EV10. Robustness of Simulation Predictions.** The sensitivity of true predictions % according to simulation to varying thresholds of ΔG_binding_) and the experimental threshold value from fluorescence experiments (x-axis). The regions within the blue and red contours are statistically significant at p values of < 0.05 and, < 0.001, respectively. The combination of thresholds within the shaded regions results in statistically significant and strong agreement wherein >65% of stable binding predictions from simulations match that of experiments.

**Figure EV11.** (A) Structural alignment of PPI complexes of FolA with different binding partners (FolD, GlyA, FolK and NudB) shows that FolA uses a similar interface to interact with multiple proteins, hence the highly significant p-value of FolA (p-value = 1.2e-9 in Fig 7B). (B) Structural alignment of PPI complexes of FolD with two different binding partners (FolA and GlyA) shows that the two interfaces are completely different, hence the non-significant p-value (p-value = 0.279 in Fig 7B).

**Figure EV12.** Z-scores for the shift in mean inter-surface contacts (observed) compared to the randomized set. The Z score measures how many standard deviations from the mean the observed value lies and determines p-value for the underlying t-statistic for the random null model (see Methods section for details). We set a significance level (e.g., 0.05), which represents the threshold for statistical significance. If the p-value is less than this significance level, we reject the null hypothesis. Otherwise, the null hypothesis is accepted.

**Figure EV13:** Overlay of representative heterodimeric structures of PPI complexes with their corresponding homodimeric structures. (A) FolA-GlyA complex superimposed with GlyA homodimer (B) PurD-PurK complex superimposed with FolK homodimer and (C) FolA-PurK complex superimposed with FolK homodimer. In all these structures, the homodimeric interface is different than the PPI interface.

**Figure EV14.** Distribution of mean pairwise inter-surface contacts for a set of 50 randomly drawn surfaces on A) FolC, and B) FolD protein. The distribution corresponds to the mean inter-surface contacts for 10000 such realizations, each with 50 randomly drawn surfaces. The blue horizontal line shows the mean value for inter-surface contacts for the random set, while the red line shows the corresponding value for the observed surfaces. A) For the FolC protein, the mean inter-surface contacts (observed) is significantly higher than the corresponding value for the random set, suggesting that the surfaces used for interaction across different proteins share a high degree of overlap. B) On the other hand, the interaction surfaces on FolD corresponding to different interaction partners shows no such overlap.

**Figure EV15.** The histogram (orange) represents a simulated population of natural enzyme k_cat_/K_M_ values drawn from a log-normal distribution centered around 10^5^ M ¹s ¹ with a spread of ±1 log unit, based on empirical data from biochemical databases. Blue vertical lines indicate the 10 representative enzyme values selected for the 10-step pathway modeled in the coarse-grained simulation. These values span the realistic catalytic efficiency range observed in natural enzymes, from 10^3^ to 10^7^ M ¹s ¹. The distribution of k_cat_/K_M_ (orange histograms) are based on data from (32).

**Figure EV16.** Progression of the 10 reactions in the diffusion-reaction model during simulation for A) monomeric enzymes where all reactions occur in the bulk, and B) when enzymes interact via an PPI-map based on experiments.

**Figure EV17.** Stoichiometry of the largest cluster at equilibrium. Imposing the folate pathway PPI in the Langevin dynamics simulations results in clusters that contain several copies of each enzyme.

**xTable EV1:** Symmetric matrix of interactions generated by taking the maximum of the fluorescence intensity generated from the two configurations. Red boxes indicate MFI > 1100, yellow for 900 < MFI < 1100, blue for 750 < MFI < 900 while white boxes indicate MFI < 750.

**Table EV2:** Mean fluorescence intensities observed for each pair of PPI tested, in both configurations. Red boxes indicate MFI > 1100, yellow for 900 < MFI < 1100, blue for 750 < MFI < 900 while white boxes indicate MFI < 750.

**Table EV3:** TM-scores of all PPI pairs.

**Table EV4:** Sequences of primers used to clone the genes corresponding to all pathway proteins explored in this study.

## Methods

### Cloning and transformation of split YFP plasmids

The genes corresponding to 41 proteins in the folate pathway were amplified from the *E. coli* genome using primers containing BamHI and XbaI sites (Table EV4). The genes were cloned in two different formats in two mutually compatible plasmids: as a fusion to the N-terminus of YFP under the control of IPTG inducible tac promoter, and as a fusion to the C-terminus of YFP under control of an arabinose inducible pBAD promoter. The YFP used in this work was the monomeric version of enhanced YFP (EYFP, truncated at 155/156) (11). This is brighter and matures faster than regular YFP, but still requires incubation at 30C for full maturation.

Out of the 41 proteins, 35 proteins were successfully cloned as fusions. NYFP fusion plasmids were transformed into BW27783 cells by TSS method (33). This method enables transformation by simple incubation at 4C without any heat shock, and thereby allows high-throughput transformation in multi-well plates. Subsequently, cells transformed with NYFP-fusion plasmids were grown and were made competent by the TSS method. For NYFP-fusion plasmid, 35 CYFP- fusion plasmids were transformed, resulting in a 35x35 matrix of all possible combinations.

### Flow cytometry to detect interaction by split YFP assay

For FACS analysis, BW27783 cells transformed with both NYFP and CYFP fusion proteins were inoculated from glycerol stocks into 200ul of supplemented M9 medium (containing Ampicillin and Chloramphenicol) in deep well plates and grown at 37C overnight. Next day, the cultures were diluted 1:100 in 150ul of supplemented M9 medium containing antibiotics and inducers (0.2% arabinose and 1mM IPTG), grown at 30C for 4-5 hours, following which YFP fluorescence was detected by FACS. For analysis, the BD LSR Fortessa was used along with the HTS (high throughput) module. For each well of the 96 well plate, 30,000 cells were analyzed, and the mean fluorescence intensity was calculated. Those wells for which less than 10,000 cells could be analyzed were discarded.

### Western blot

To detect intracellular abundance of YFP fusion proteins, BW27783 cells transformed with a single plasmid (either NYFP or CYFP fusion proteins) were used. Cultures were inoculated from glycerol stocks into 200ul of supplemented M9 medium (containing Ampicillin or Chloramphenicol, depending on the plasmid) in deep well plates and grown at 37C overnight. Next day, the cultures were diluted 1:100 in 5ml supplemented M9 medium containing antibiotics and inducers (0.2% arabinose or 1mM IPTG) in a 24-well deep well plate and grown at 30C for 4-5 hours. The cultures were spun down, and lysis was carried out in 100ul of Bugbuster solution in 1xTBS for 30 minutes while shaking. Complete EDTA free protease inhibitor cocktail from Roche was added during the lysis step. Subsequently, 100ul lysate was mixed with 20ul of 50% glycerol+10% SDS solution and heated at 95°C for 10 mins. This dissolves the lysate and results in a clear solution. 15ul of the cleared lysate was mixed with 5ul of 4x loading dye, heated at 95C for 15 minutes and loaded on to a 4-12% Bis-tris gel. The left- over sample was used for estimation of total cellular protein content using a BCA kit.

Western breeze chromogenic immuno-detection kit (Thermo) was used for western blot, following manufacturer’s guidelines. For detection of YFP fusion proteins, polyclonal anti-GFP antibody (Cat # A11122 from Invitrogen) was used as a primary antibody at a dilution of 1:2000, while goat anti-rabbit antibody conjugated to alkaline phosphatase was used as a secondary antibody. ImageJ was used to quantify the bands following blotting, and the intensities were normalized by total protein abundance obtained from BCA method.

### Microscopy

BW27783 cells transformed with both NYFP and CYFP fusion proteins were inoculated from glycerol stocks into 2ml of supplemented M9 medium (containing Ampicillin and Chloramphenicol) and grown at 37C overnight. Next day, the cultures were diluted 1:100 in 5ml of supplemented M9 medium containing antibiotics and inducers (0.2% arabinose and 1mM IPTG), grown at 30C for 4-5 hours. For live phase contrast images, 2ul of the culture was directly spotted on 1.5% low melting agarose (Calbiochem) pads. The agarose was dissolved in supplemented M9 medium. Pads were then flipped on class #1.5 glass dish (Willco Wells), and the images were acquired at room temperature with Zeiss Cell Observer microscope.

### Library generation

Based on the structure of DHFR (PDB: 7DFR), we selected all surface exposed residues of DHFR (accessibility > 80%), excluding its active site and those contacting active site residues within 5A radius. There were 85 such positions in DHFR that satisfied the above criteria, and at each of these positions, a VDS library was created. VDS codes for a combination of [ACG][AGT][GC] bases, which in total code for 1530 codons and 1105 amino acids. These codons code for all different amino acid types (hydrophobic, polar, charged), however does not incorporate large hydrophobics (F, W or Y) or proline, and does not incorporate any stop codon. The mutagenesis was carried out on the NYFP-DHFR plasmid in a 96-well PCR plate using a megaprimer based method, following which the reactions from individual wells were combined, digested overnight using DpnI to remove the WT plasmid and transformed into DH5-alpha competent cells. A small amount was plated for single colonies, while the rest of the culture was grown overnight in the presence of antibiotics. The plasmid isolated from single clones were sequenced to confirm presence of mutations. Following this, the library plasmid was isolated from the culture using miniprep.

### Sorting of clones

The NYFP-DHFR VDS library plasmid pool was transformed into the background of BW27783 cells containing CYFP-MetF/CYFP-FolK plasmids using a cell: plasmid ratio of 100:1 to minimize chances of multiple plasmids transformed into a single cell. The cultures were grown overnight at 37C, and next day they were diluted 1:100 into fresh supplemented M9 medium containing antibiotics and inducers (0.2% arabinose and 1mM IPTG). Using the BD FACS Aria Cell Sorter, single clones from the top and bottom 5% of the fluorescence intensity distribution of both cultures were sorted into wells of a 96-well plate that contained fresh supplemented M9 medium without any antibiotics. The single clones were allowed to grow for 3 days, following which they were diluted in medium containing antibiotics and allowed to grow overnight. Next day, after growth in the presence of inducers, fluorescence intensity was measured on the Fortessa analyzer, and those that had substantial change in fluorescence compared to the WT culture were sent for sequencing to identify the mutation.

### Permutation-based p-value estimation for statistical significance of PPI clusters

To assess the statistical significance of observed interaction strengths within specified protein clusters, we employed a non-parametric permutation test. For each cluster, we computed a summary statistic (in this case, median) of the observed fluorescence values and compared it to a null distribution generated by randomly sampling an equal number of protein pairs from the full dataset. This process was repeated 10,000 times to build the empirical null distribution. The empirical p-value was calculated as the proportion of null statistics greater than or equal to the observed value, using the following formula:

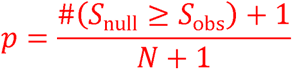

where S_obs_ is the observed statistic, S_null_ are the null statistics, and N is the number of permutations. This method of calculating p-value does not require any assumption of normal distribution of the data.

### Raman Spectroscopy

To better understand any plausible interaction between DHFR and FolK we performed Raman spectroscopy. As a vibrational spectroscopy technique Raman spectroscopy provides a host of structural information and hence has been in use for mechanistically probing conformational changes in proteins (34–36). As compared to other spectroscopic techniques Raman spectroscopy is essentially a non-destructive technique and is known for its high sensitivity. Raman spectroscopy is an informative technique in probing conformational changes upon binding interactions in protein (37). Conformational changes primarily impact the Amide I range (1600 to 1700 cm^-1^) which arise due to C=O stretching vibrations of the peptide bonds in turn modulated by the secondary structural elements viz. helices, sheets, turns etc. We probed into the secondary structure of DHFR and FolK using Raman spectroscopy with an aim to investigate the solution status after mixing. Any interaction upon mixing would essentially give rise to new spectral features which upon careful examination help in mechanistic understanding of conformational changes upon binding. Control experiments were run in presence of ADK (detailed in the results section) and protein concentration in all the acquisition was kept at 15 µM.

The Raman spectroscopy studies utilized a Horiba XploRa confocal Raman microscope, featuring a thermoelectrically cooled detector at -70°C and a 1200 gr/mm grating optimized for 750nm. A 785 nm solid-state laser was employed as the excitation source, with each measurement lasting 180 seconds. To ensure accuracy, four spectra were gathered for each sample, facilitating cosmic ray removal. Spectral smoothing was achieved using the Horiba denoise algorithm, and a polynomial fit was applied for fluorescent baseline correction prior to peak deconvolution in Labspec 6. The 785 nm laser’s power was maintained at ∼41 mW to avoid photo-bleaching or thermal damage, given the excitation wavelength’s significant separation from protein absorption bands. Settings included a 200 µm spectrometer slit, a 500 µm confocal aperture, and calibration against a 520.7 cm^-1^ silicon reference. Experiments used 20 µM protein solutions in sodium phosphate buffer, with a 20µl volume, alongside buffer blanks for thorough comparison. Spectral data were normalized against the distinct 330 cm^-1^ peak.

### Protocol for generating random surfaces

To ensure that the difference in inter-surface contacts for the observed and randomly generated surfaces are not merely due to differences in the surface sizes, we followed the following protocol while generating random surfaces.

For each of the 10 folate pathway proteins under study, we compute the radius of gyration of the residues that make up the ‘n’ interaction surfaces for each protein. The mean radius of gyration for the surface is inferred from these values. This is used as the radius of an idealized spherical interaction surface.

1. To draw the random surfaces, we define a list of solvent accessible residues for every folate protein based on DSSP calculations. All residues with relative solvent exposed surface area > 0.25 were considered exposed and were considered while defining random surfaces. For every random surface, we pick a residue from this exposed residue list and draw a probe radius (computed in step (i)) around it to define a set of exposed residues that make up the random surface.
2. To account for the irregularities in protein surfaces, and the accompanying variability in the number of C_α_ atoms that make up each surface, we employ a normalized contact number to compute statistical differences. The raw contacts are normalized by m_i_*m_j,_ where mi and mj refer to the number of C atoms that are part of the i^th^ and j^th^ random surfaces (see Eq. 2)

### Metadynamics Simulations

35 enzymes corresponding to the Folate, Purine and Pyrimidine pathways were used in the computational study. These correspond to the same set of genes listed in the experimental PPI matrix shown in Fig 2. We use AlphaFold 2.0 multimer build to predict the pairwise dimeric structures for all intra-pathway pairs within the 3 pathways. We also randomly sample 55 dimers each corresponding to the Folate-Purine and Purine-Pyrimidine interactions to estimate inter-pathway PPIs.

To account for dynamicity in interactions and compute the stability of the AlphaFold predicted dimers, we employed metadynamics simulations wherein the distance between two chains in the dimer is used as the collective variable of interest (Fig 5A). We then compute the free energy landscape as a function of these inter-chain distances for every folate dimer under study (Fig 5A).

Minima at shorter distances (< interaction radii of the pair) corresponds to a favorable dimer. We then integrated over all bound and unbound structures in the landscape to compute the binding free energy (ΔG_binding_ in Fig 5B). All simulations were performed using NAMD2.3 (38) with the CHARMM36 forcefield. Interaction cutoffs of 12 A and 13.5 A were set for van der Waals and electrostatic interactions. A simulation timestep of 2 fs was used for the study. Meta dynamics simulations were run till the PMFs exhibited convergence.

### Statistical test to check for non-random overlap of interaction surfaces

To check if there is a non-random overlap of interaction surfaces across different interaction partners, we define an order parameter -- inter-surface Cα- Cα contacts as a measure of shared interaction surfaces. To compute this quantity, we define for each folate protein an interaction surface --a list of interacting residues -- with every other protein that it interacts with in the folate pathway. To draw this residue list, we use structures that populate the free energy minima in our metadynamics simulations or revealed by high throughput mutagenesis experiment. For instance, in metadynamics simulations FolA shows stable interactions with 4 other partners. Therefore, FolA has 4 potentially distinct interaction surfaces for each of these different partners. We then compute Ca-Ca contacts for the set of FolA residues that define each of the 4 surfaces (across all possible surface pairs) and then compute a quantity -- *MISC^O^* - Mean Inter-surface Contacts derived from actual PPI surfaces obtained from mutational mapping or metadynamics simulations -- for each of the 10 folate proteins.

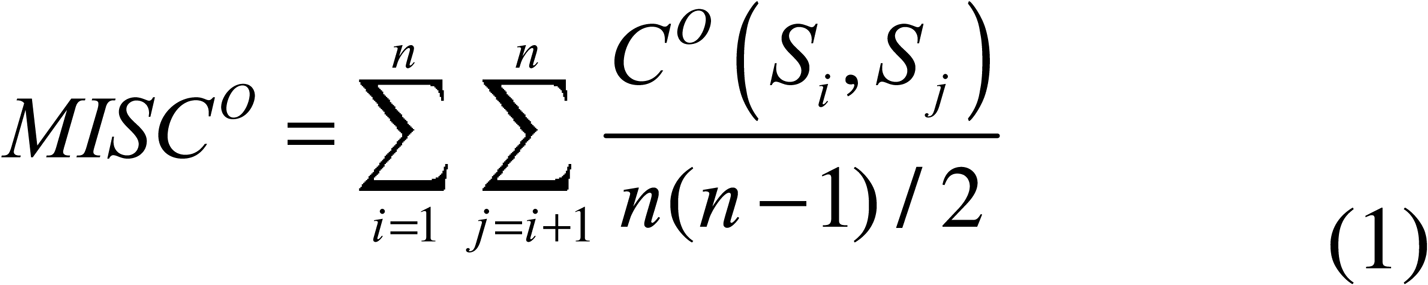

Here superscript *O* refers to observed (as opposed to random control) surfaces. *n* is the number of detected interaction surfaces for the protein. *S_i_* and *S _j_* represent the list of residues that make up the *i* and *j*’th surfaces, respectively. The function *C^O^* (*S*, *S*) represents the normalized number of pairwise C_α-_ C_α_ contacts between residues that make up the *i* and *j*th surfaces.

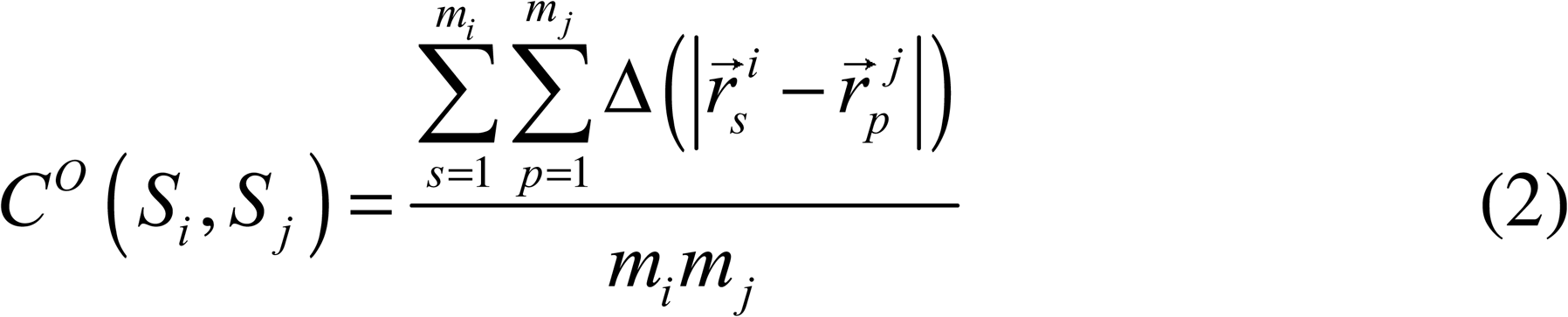

Here, *m_i_*is the number of residues belonging to *i*-th surface, 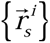 represents set of coordinates of all *C_α_* atoms belonging to surface *i* and

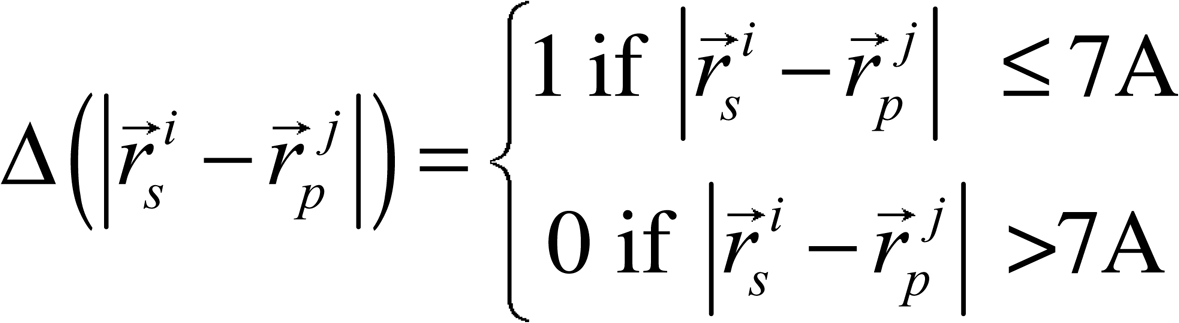

Is a contact function that defines contact between *C_α_* atoms belonging to different surfaces with 7A cutoff.

If the interaction surfaces for different partners exactly overlap, the mean inter-surface contacts (observed) would be high. On the other hand, if these surfaces were distinct and non-overlapping, this quantity would be very small. We compute these mean inter-surface contacts (observed) for each of the 10 enzymes of the folate pathway.

To determine the statistical significance of the overlap between different interacting surfaces we propose a null hypothesis that the interaction surfaces are randomly distributed on the surface of a protein in question. In other words, we test whether the mean inter-surface contacts (observed) are significantly higher than the analogous quantity *MISC^R^* – Mean Inter-surface Contacts (Random) -- for a set of 100 randomly drawn surfaces for every folate protein under study.

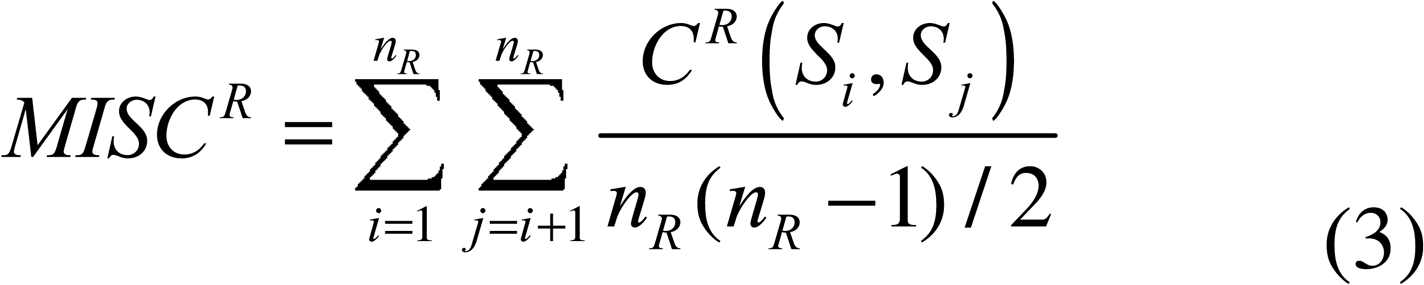

Where *n_R_* = 50 the number of randomly generated surfaces The function *C^R^* (*S_i_*, *S_j_*) represents the normalized number of pair-wise C_α-_ C_α_ contacts between the i and jth randomly generated surfaces, defined analogously to Eq.2 but using *C_α_* coordinates of randomly generated surfaces (see below for the algorithm that generated random surfaces). The mean inter-surface contacts (random) are then compared to the inter-surface contacts observed in experiment and simulations to assess the non-random nature of their overlap. 10,000 such random sets are generated, and the distribution of the inter-surface contacts in the random control is then computed (Extended View Fig EV14). Our null hypothesis is that there is no significant difference between the observed mean normalized contacts and what would be expected by random chance. In other words, any observed difference is due to random variation per the null hypothesis.

To determine the significance of the observed test statistic, we perform a t-test for statistical significance. Here, the Z score is calculated using the following equation,

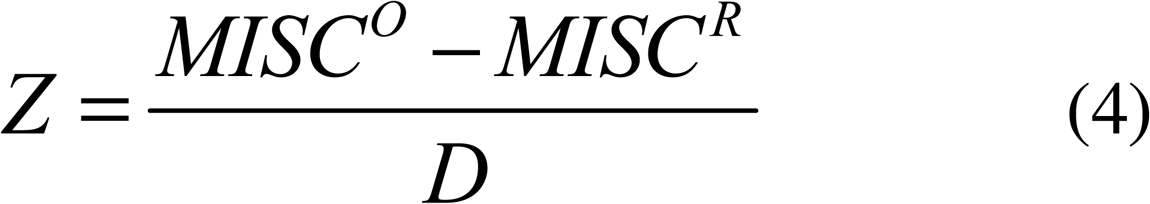

Here, the denominator *D* refers to the variation in the inter-surface contacts, for the randomly generated surfaces.

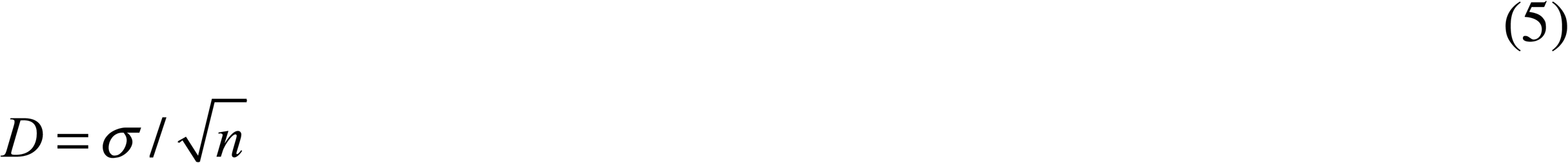

Here, *σ* refers to the standard deviation in the normalized pair-wise contact number (equation 2) for the randomly generated surfaces whereas ‘n’ refers to the number of randomly generated sets used for the computation of the statistic. The Z score measures how many standard deviations from the mean the observed value lies and determines p-value for the underlying t-statistic for the random null model We set a significance level (e.g., 0.05), which represents the threshold for statistical significance. If the p-value is less than this significance level, we reject the null hypothesis. Otherwise, the null hypothesis is accepted.

### Langevin Dynamics (LD) Simulations

#### The Patchy Particle Enzyme Model

To study how protein-protein interactions between enzymes modulates reaction fluxes in the folate biosynthetic pathway, we employed off-lattice Langevin dynamics (LD) simulations implemented in the LAMMPS (39) molecular dynamics engine. This approach captures both thermal motion and viscous damping of particles, allowing us to simulate the coupled effects of diffusion, interaction, and reaction in a biologically realistic environment. Patchy hard-sphere models are widely used to study the self-assembly of multivalent proteins (40, 41). In our framework, each enzyme is represented as a rigid hard sphere bearing two adhesive patches (Fig 8A) that mimic -- i) catalytic active site and, ii) interface for protein protein interaction analogous to the interfaces in Fig 7. The catalytic patches carry distinct “identities,” ensuring that only one specific ligand type can bind to each active site. Similarly, the protein-protein interaction interface patch also have 10 distinct identities corresponding to 10 different enzymes in the Folate pathway.

Substrate molecules themselves are modeled as diffusing hard spheres which react only upon meeting their complementary patch. In Fig 8A, the enzyme core is shown as a larger gray sphere, while its active-site patches and protein-protein interaction patches are shown as smaller colored hemispheres on the surface of the large spheres. Although substrate molecules may transiently associate anywhere on the enzyme surface via these non-specific forces, a catalytic event occurs only when a ligand contacts its matching active-site patch.

We model a sequential enzymatic pathway consisting of N turnover steps (see Eqn. 6 and Fig 1A). In this scheme, each enzyme catalyzes a transformation in the sequence

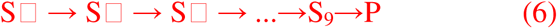

so that, after N consecutive reactions, the initial substrate **S_0_** is converted into the final product **P** within an enzyme cluster or in the bulk.

In these simulations, the enzyme particles are surrounded by a pool of substrates which are entirely in an S0 state corresponding to the first reaction in the pathway. To capture both diffusion and catalysis, we implement a stochastic diffusion-reaction algorithm: each time a substrate or intermediate S_i_ occupies its cognate active site patch on enzyme E_i_, it may proceed to S_i+1_ with probability Preact during that timestep. Whenever a substrate or intermediate binds to its matching active site patch, it may convert to the next chemical state with probability P_react_. Each active site patch supports two classes of binding:

- Cognate binding (strength ε_co_) occurs only between an active site patch and its specific ligand, ensuring that each enzyme catalyzes a defined step in the linear pathway (Eqn. 1). The strength of cognate binding in our simulations is 5 kBT.
- Non cognate binding (strength ε_nc_ = ε_PL_) represents all other, non-specific associations between patches and off target substrates or intermediates. The strength of non-cognate binding in our simulations is 0.8 k_B_T. In our model, ε_nc_ is set equal to the generic enzyme–ligand interaction ε_PL_, so that any non-complementary patch–ligand pair interacts solely via the same weak, non-specific forces governing the enzyme surface and ligand molecules.

### Potential Functions and Simulation Parameters

All non specific interactions—enzyme–enzyme interactions outside the PPI patch (ε_PP_), enzyme–ligand outside the active site (ε_PL_), and non cognate patch–ligand contacts (ε_nc_)—are described by a standard Lennard–Jones potential.

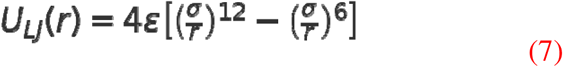

where σ is the sum of the two particle radii (σenz = 20 Å for enzyme cores; σpatch = σligand = 8 Å for patches and ligands), and the cutoff distance rc = 2.5σ. The depth ε of the attractive well is chosen to set the desired interaction strength— ε_PP_ for protein–protein, ε_PL_ for non specific protein–ligand, and ε_nc_ for non cognate binding are set to 0.8 kT.

Specific (cognate) binding between a) an active site patch and its matching ligand, and b) protein-protein interaction patches on enzymes is implemented via a Morse potential such that,

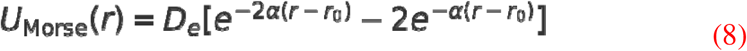

Here, D_e_ controls the strength of the attractive interactions, r0 is the equilibrium distance for the interaction and α is a parameter that controls the sharpness of the potential. For protein-protein interactions, the interaction patches on enzymes experience short-range attractive interactions, with α = 2.2 Å ¹, and r = 17.056 Å. The cutoff distance was set to 20 Å. The range of the attractive interaction between protein-protein interaction patches in our simulations is set to 2.8 Å enforcing single valent binding per patch. The strength of interaction between any two PPI-patches is modeled based on the fluorescence intensities of the corresponding enzyme pair in experiments (Fig 2A). The K_D_ values for all interacting enzyme pairs with MFI > 1000 is set to 10μM. For MFI between 800-1000, the K_D_ for the interaction is set to 200 μM. For MFI between 650-800, the K_D_ is set to 500 μM. All MFI < 650 are modeled with a K_D_ of 100 mM. These K_D_ are translated into interaction strengths between protein-protein interaction patches in the Morse-potential such that, D_e_ = K_B_T.ln(K_D_). For sake of computational simplicity, we assume that the entropic contribution to the free energy is negligible.

This interaction facilitates transient enzyme clustering, enhancing local concentration. Similarly, cognate binding of substrate onto the complementary active site on enzymes is modeled via the Morse potential with α = 1 Å ¹, and r = 12.7 Å. The range of the attractive interaction between substrate and a cognate binding site is 2.5 Å ensuring that only one substrate molecule can bind to an active site at any given time. For cognate binding, D_e_= ε_co_ = 5 kT.

The equations of motion for each rigid patchy particle is

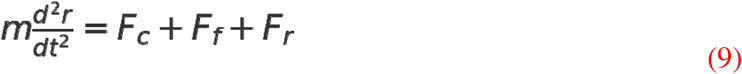

where Fc= -delta_U is the sum of conservative forces from Eqs. 7 and 8.

Ff = -α v is the viscous drag, and Fr is a stochastic force with

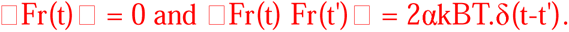

Simulations were carried out in LAMMPS under NVT conditions using a Langevin thermostat at T = 300 K and solvent viscosity η = 10 ³ Pa·s.

The damping coefficient α governs how rapidly momentum is relaxed and is related to solvent viscosity and particle radius by:

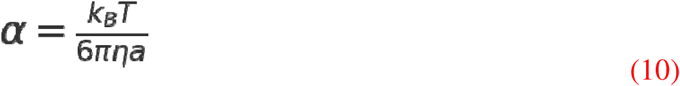

We employed a timestep Δt = 30 fs. Because each enzyme is treated as a rigid multi center body (no internal degrees of freedom), we used LAMMPS’s rigid particle integrator to enforce fixed relative positions of the core and its patches.

The simulation box, in our study, contains 300 copies of each of the 10 coarse-grained enzyme types such that the effective concentrations of enzymes in the simulation box is 50 uM. In order to facilitate collisions between enzyme molecules at the simulation timescale, during the first 1 us of the simulation the enzyme molecules are confined within a spherical region of 400 Å radius in the center of the cubic simulation box. The spherical confinement is then relaxed, and th enzyme clusters are allowed to equilibrate during the rest of the simulation run.

### Parameterization of Reaction Probabilities

To model enzymatic turnover in a coarse-grained simulation of a 10-step metabolic pathway, we derived per-attempt reaction probabilities for each enzymatic step based on the known distribution of k_cat_/K_M_ values observed in natural enzymes. A total of 10 representative kcat/KM values were randomly sampled from a log-normal distribution with parameters derived from the enzyme k_cat_/K_M_ distribution presented in Extended View Fig EV15 (based on data by (32), which spans approximately seven orders of magnitude (10^2^ to 10^9^ M^-1^s^-1^). We used a normal distribution in log-space with a mean of 5 and standard deviation of 1.0 to generate log_10_(k_cat_/K_M_) values. These were exponentiated to yield real-world k_cat_/K_M_ values. The value of P_react_ for the 10 rxns is chosen based on a known distribution of k_cat_/K_M_ values (normalized by the k_cat_/K_M_ for the diffusion limit → 10^8^ M^-1^s^-1^) (see Methods and Extended View Fig EV15). Each sampled value was then normalized by the diffusion limit, taken to be 10^8^ M^-1^s^-1^, which reflects the maximum catalytic efficiency constrained by molecular diffusion in the cytoplasm. The resulting normalized reaction efficiencies, η_react_ = (k_cat_/K_M_)/10^8^ represent the relative catalytic efficiency of each enzyme compared to an idealized diffusion-limited enzyme. Note that η_react_ is a dimensionless quantity which we then use to set values of P_react_ for the 10 reactions in our diffusion-reaction model (see Fig EV15).

### Data availability

This study includes no data deposited in external repositories.

## References

1. P. A. Srere, Complexes of sequential metabolic enzymes. Annu Rev Biochem 56, 89–124 (1987).

2. J. B. Robinson, Jr., P. A. Srere, Organization of Krebs tricarboxylic acid cycle enzymes. Biochem Med 33, 149–157 (1985).

3. S. An, R. Kumar, E. D. Sheets, S. J. Benkovic, Reversible compartmentalization of de novo purine biosynthetic complexes in living cells. Science 320, 103–106 (2008).

4. Y. Deng et al., Mapping protein-protein proximity in the purinosome. J Biol Chem 287, 36201–36207 (2012).

5. J. B. French et al., Spatial colocalization and functional link of purinosomes with mitochondria. Science 351, 733–737 (2016).

6. L. J. Sweetlove, A. R. Fernie, The role of dynamic enzyme assemblies and substrate channelling in metabolic regulation. Nat Commun 9, 2136 (2018).

7. Y. Zhang, A. R. Fernie, Metabolons, enzyme-enzyme assemblies that mediate substrate channeling, and their roles in plant metabolism. Plant Commun 2, 100081 (2021).

8. F. Wu, S. Minteer, Krebs cycle metabolon: structural evidence of substrate channeling revealed by cross-linking and mass spectrometry. Angew Chem Int Ed Engl 54, 1851–1854 (2015).

9. S. Bhattacharyya et al., Transient protein-protein interactions perturb E. coli metabolome and cause gene dosage toxicity. Elife 5, e20309 (2016).

10. S. Bhattacharyya, S. Bershtein, B. V. Adkar, J. Woodard, E. I. Shakhnovich, Metabolic response to point mutations reveals principles of modulation of in vivo enzyme activity and phenotype. Mol Syst Biol 17, e10200 (2021).

11. K. Ohashi, T. Kiuchi, K. Shoji, K. Sampei, K. Mizuno, Visualization of cofilin-actin and Ras-Raf interactions by bimolecular fluorescence complementation assays using a new pair of split Venus fragments. Biotechniques 52, 45–50 (2012).

12. S. Bershtein, W. Mu, A. W. Serohijos, J. Zhou, E. I. Shakhnovich, Protein quality control acts on folding intermediates to shape the effects of mutations on organismal fitness. Mol Cell 49, 133–144 (2013).

13. D. B. Lukatsky, B. E. Shakhnovich, J. Mintseris, E. I. Shakhnovich, Structural similarity enhances interaction propensity of proteins. J Mol Biol 365, 1596–1606 (2007).

14. Y. Zhang, J. Skolnick, TM-align: a protein structure alignment algorithm based on the TM-score. Nucleic Acids Res 33, 2302–2309 (2005).

15. T. J. Kappock, S. E. Ealick, J. Stubbe, Modular evolution of the purine biosynthetic pathway. Curr Opin Chem Biol 4, 567–572 (2000).

16. Y. Zhang, M. Morar, S. E. Ealick, Structural biology of the purine biosynthetic pathway. Cell Mol Life Sci 65, 3699–3724 (2008).

17. W. Gallagher, FTIR analysis of protein structure. Course manual Chem 455 (2009).

18. S. Ranganathan, J. Liu, E. Shakhnovich, Enzymatic metabolons dramatically enhance metabolic fluxes of low-efficiency biochemical reactions. Biophys J 122, 4555–4566 (2023).

19. P. J. Stover, M. S. Field, Trafficking of intracellular folates. Adv Nutr 2, 325–331 (2011).

20. Y. Zheng, L. C. Cantley, Toward a better understanding of folate metabolism in health and disease. J Exp Med 216, 253–266 (2019).

21. C. L. Krumdieck, K. Fukushima, T. Fukushima, T. Shiota, C. E. Butterworth, Jr., A long-term study of the excretion of folate and pterins in a human subject after ingestion of 14C folic acid, with observations on the effect of diphenylhydantoin administration. Am J Clin Nutr 31, 88–93 (1978).

22. A. E. von der Porten et al., In vivo folate kinetics during chronic supplementation of human subjects with deuterium-labeled folic acid. J Nutr 122, 1293–1299 (1992).

23. V. Schirch, W. B. Strong, Interaction of folylpolyglutamates with enzymes in one-carbon metabolism. Arch Biochem Biophys 269, 371–380 (1989).

24. L. Madrova et al., Mass spectrometric analysis of purine de novo biosynthesis intermediates. PLoS One 13, e0208947 (2018).

25. M. Castellana et al., Enzyme clustering accelerates processing of intermediates through metabolic channeling. Nat Biotechnol 32, 1011–1018 (2014).

26. B. Bulutoglu, K. E. Garcia, F. Wu, S. D. Minteer, S. Banta, Direct Evidence for Metabolon Formation and Substrate Channeling in Recombinant TCA Cycle Enzymes. ACS Chem Biol 11, 2847–2853 (2016).

27. I. Skalidis et al., Structural analysis of an endogenous 4-megadalton succinyl-CoA-generating metabolon. Commun Biol 6, 552 (2023).

28. O. Peleg, J. M. Choi, E. I. Shakhnovich, Evolution of specificity in protein-protein interactions. Biophys J 107, 1686–1696 (2014).

29. J. D. Han et al., Evidence for dynamically organized modularity in the yeast protein-protein interaction network. Nature 430, 88–93 (2004).

30. F. M. Meyer et al., Physical interactions between tricarboxylic acid cycle enzymes in Bacillus subtilis: evidence for a metabolon. Metab Eng 13, 18–27 (2011).

31. J. Mowbray, V. Moses, The tentative identification in Escherichia coli of a multienzyme complex with glycolytic activity. Eur J Biochem 66, 25–36 (1976).

32. A. Bar-Even, R. Milo, E. Noor, D. S. Tawfik, The Moderately Efficient Enzyme: Futile Encounters and Enzyme Floppiness. Biochemistry 54, 4969–4977 (2015).

33. C. T. Chung, S. L. Niemela, R. H. Miller, One-step preparation of competent Escherichia coli: transformation and storage of bacterial cells in the same solution. Proc Natl Acad Sci U S A 86, 2172–2175 (1989).

34. R. Tuma, Raman spectroscopy of proteins: from peptides to large assemblies. Journal of Raman Spectroscopy: An International Journal for Original Work in all Aspects of Raman Spectroscopy, Including Higher Order Processes, and also Brillouin and Rayleigh Scattering 36, 307–319 (2005).

35. A. Rygula et al., Raman spectroscopy of proteins: a review. Journal of Raman Spectroscopy 44, 1061–1076 (2013).

36. J. M. Benevides, S. A. Overman, G. J. Thomas Jr, Raman spectroscopy of proteins. Current protocols in protein science 33, 17.18. 11–17.18. 35 (2003).

37. J. Lippert, D. Tyminski, P. Desmeules, Determination of the secondary structure of proteins by laser Raman spectroscopy. Journal of the American Chemical Society 98, 7075–7080 (1976).

38. J. C. Phillips et al., Scalable molecular dynamics on CPU and GPU architectures with NAMD. J Chem Phys 153, 044130 (2020).

39. A. P. A. Thompson, H. M.; Berger, R.; Bolintineanu, D. S.; Brown, W. M.; Crozier, P. S. ; Veld, P. J. in ’t; Kohlmeyer, A; Moore, S.G.; Nguyen, T. D.; Shan, R.; Stevens, M. J.; Tranchida, J; Trott, C; Plimpton, S. J.;, LAMMPS - a flexible simulation tool for particle- based materials modeling at the atomic, meso, and continuum scales. Comp. Phys. Comm. 271 (2022).

40. J. R. Espinosa et al., Liquid network connectivity regulates the stability and composition of biomolecular condensates with many components. Proc Natl Acad Sci U S A 117, 13238–13247 (2020).

41. A. R. Tejedor, A. Garaizar, J. Ramirez, J. R. Espinosa, ’RNA modulation of transport properties and stability in phase-separated condensates. Biophys J 120, 5169–5186 (2021).

