## Supplementary figures and images for "Conserved interfaces mediate multiple protein-protein interactions in a prokaryotic metabolon"

### Supplementary Fig 1

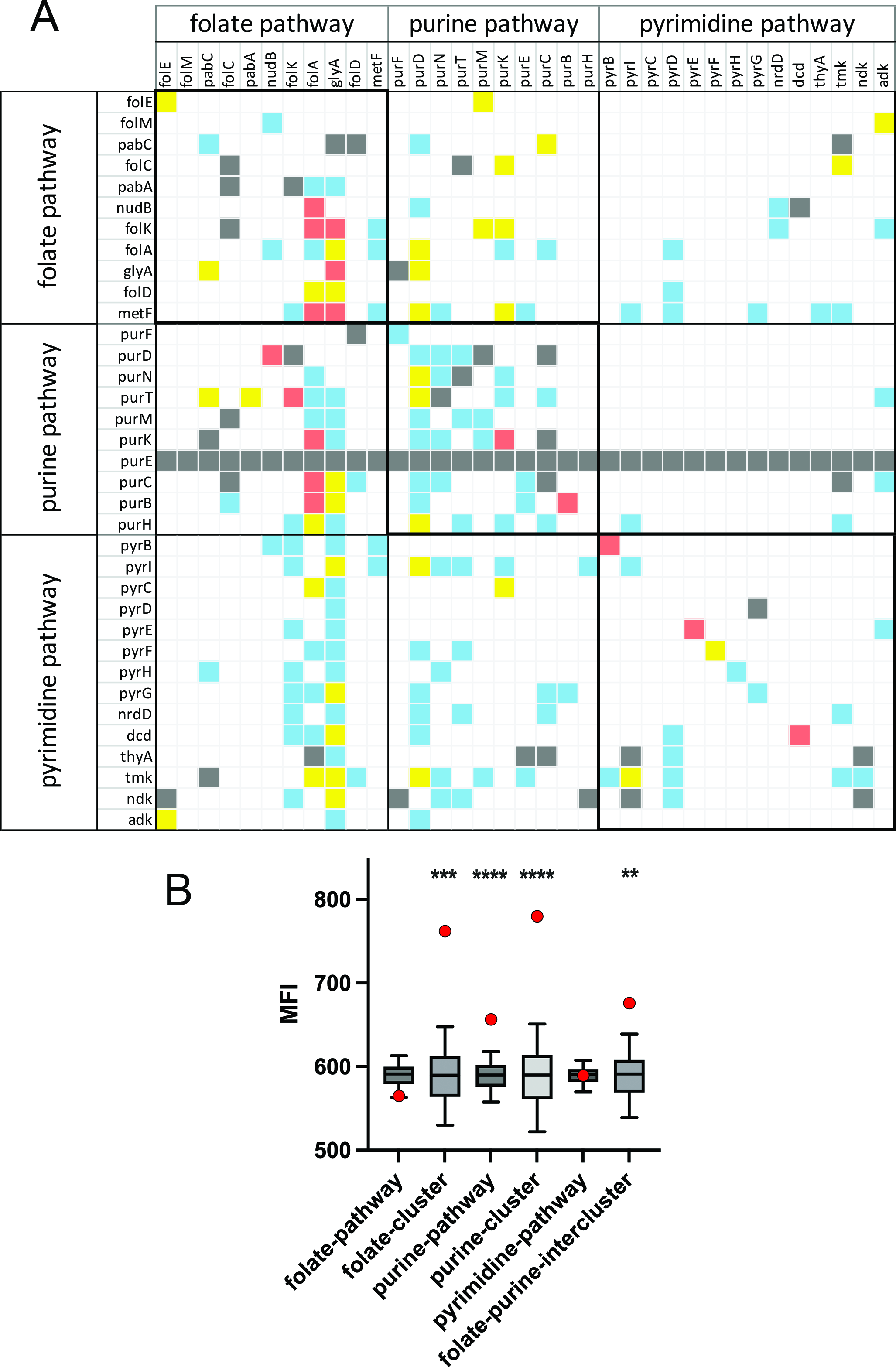

### Supplementary Fig 2

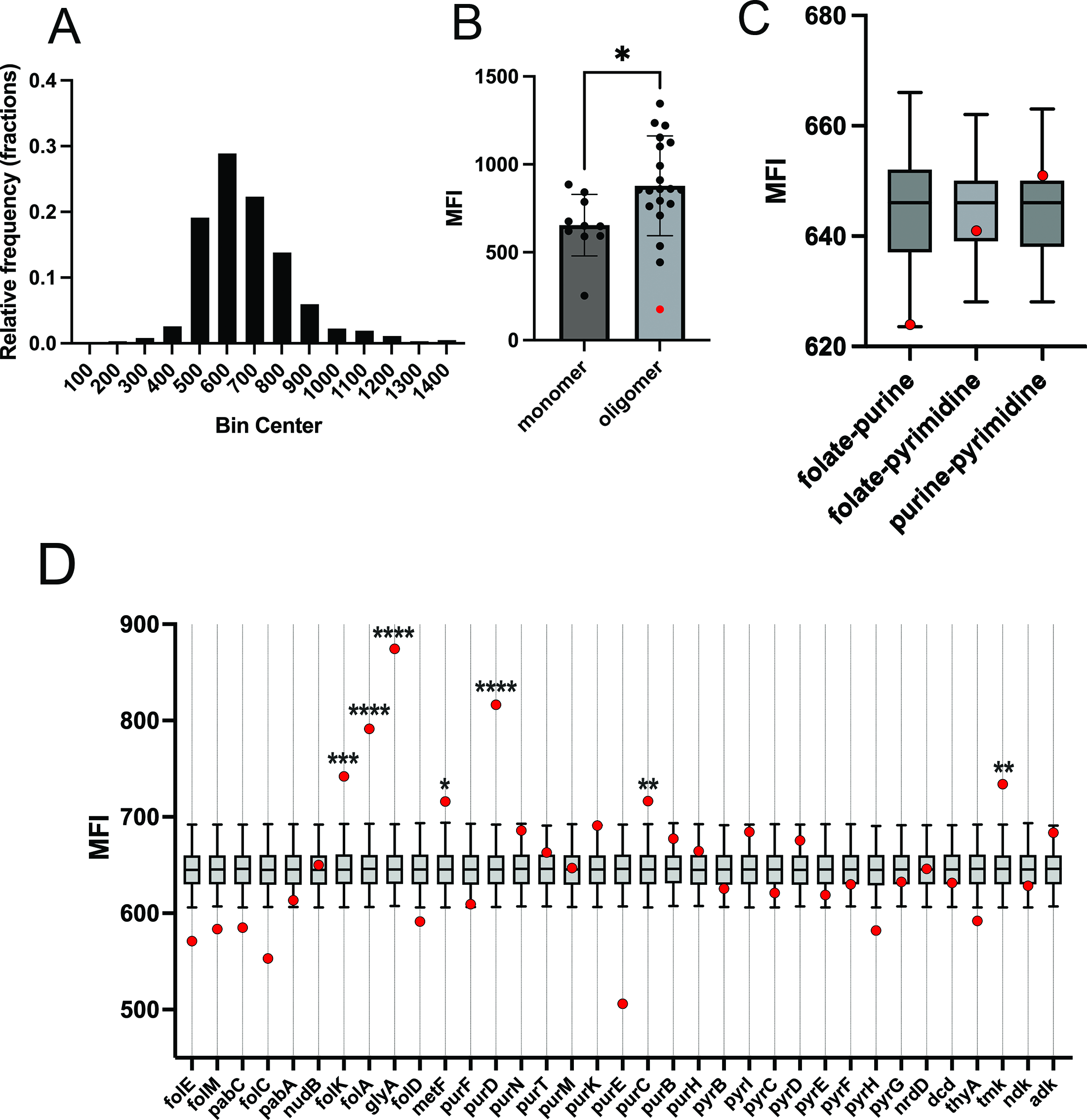

### Supplementary Fig 3

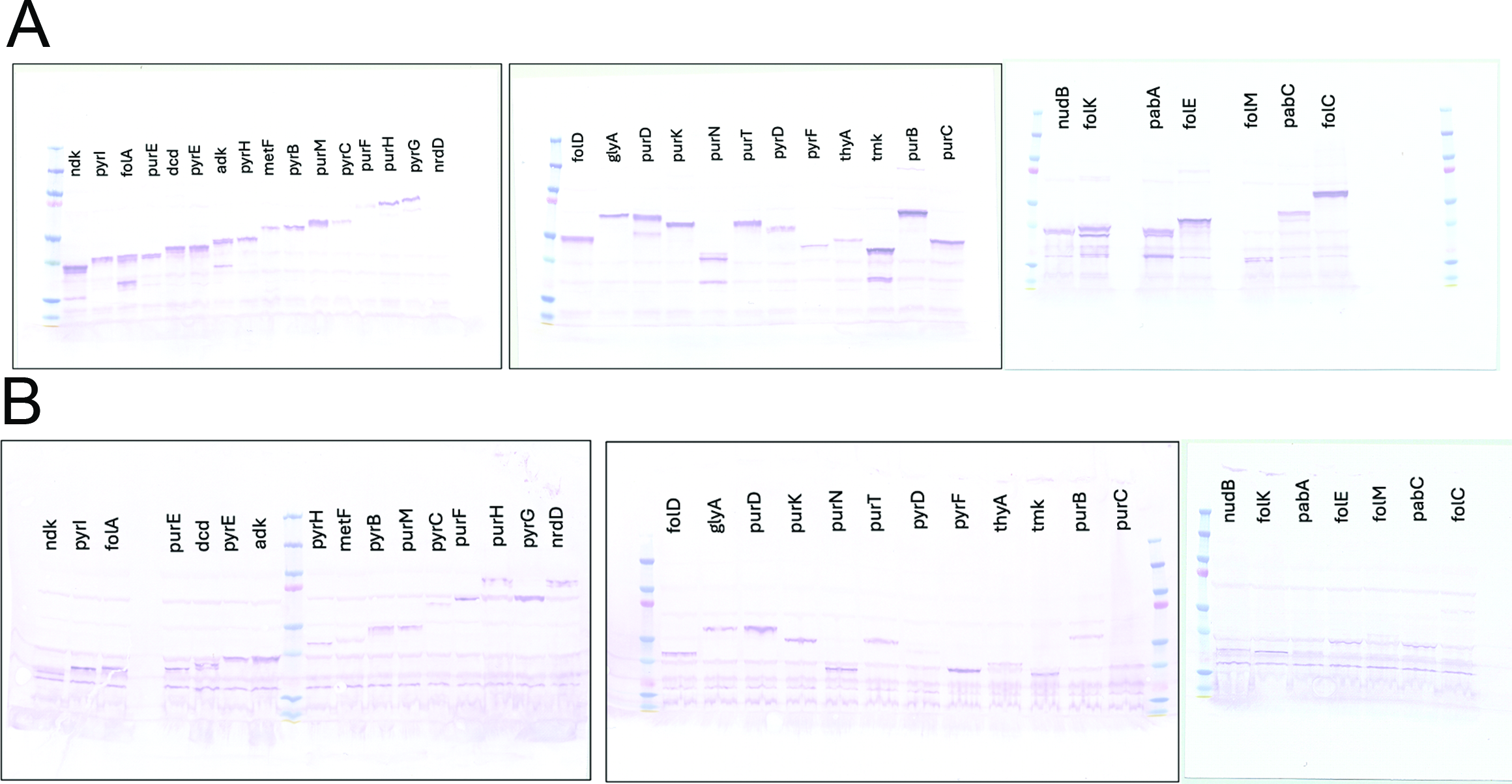

### Supplementary Fig 4

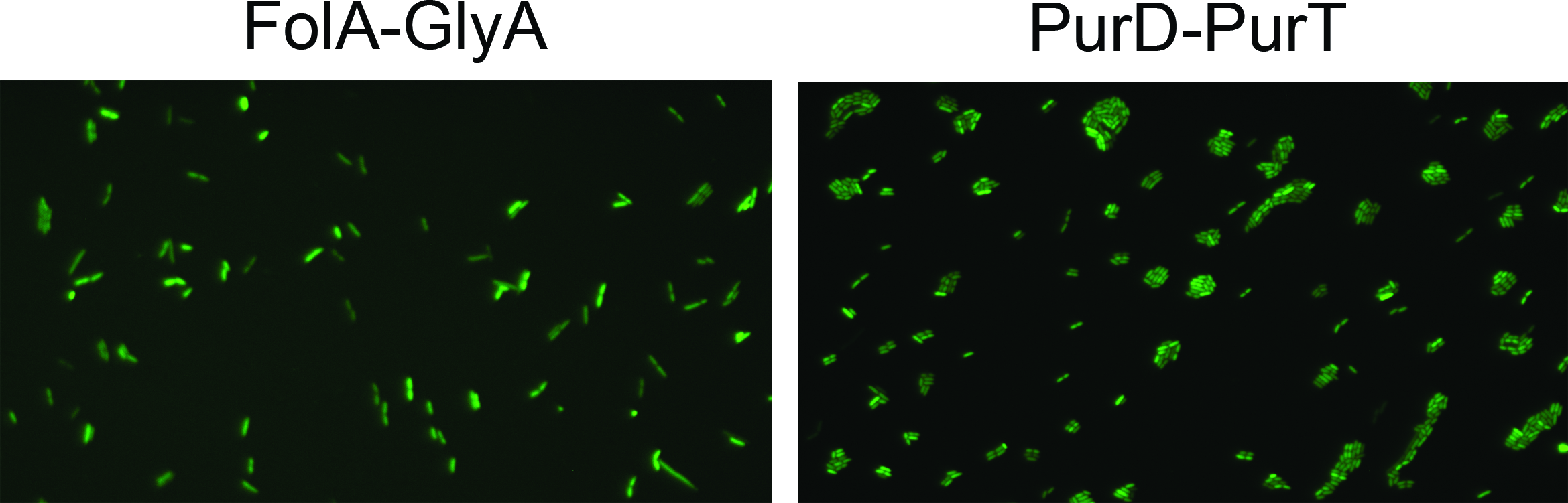

### Supplementary Fig 5

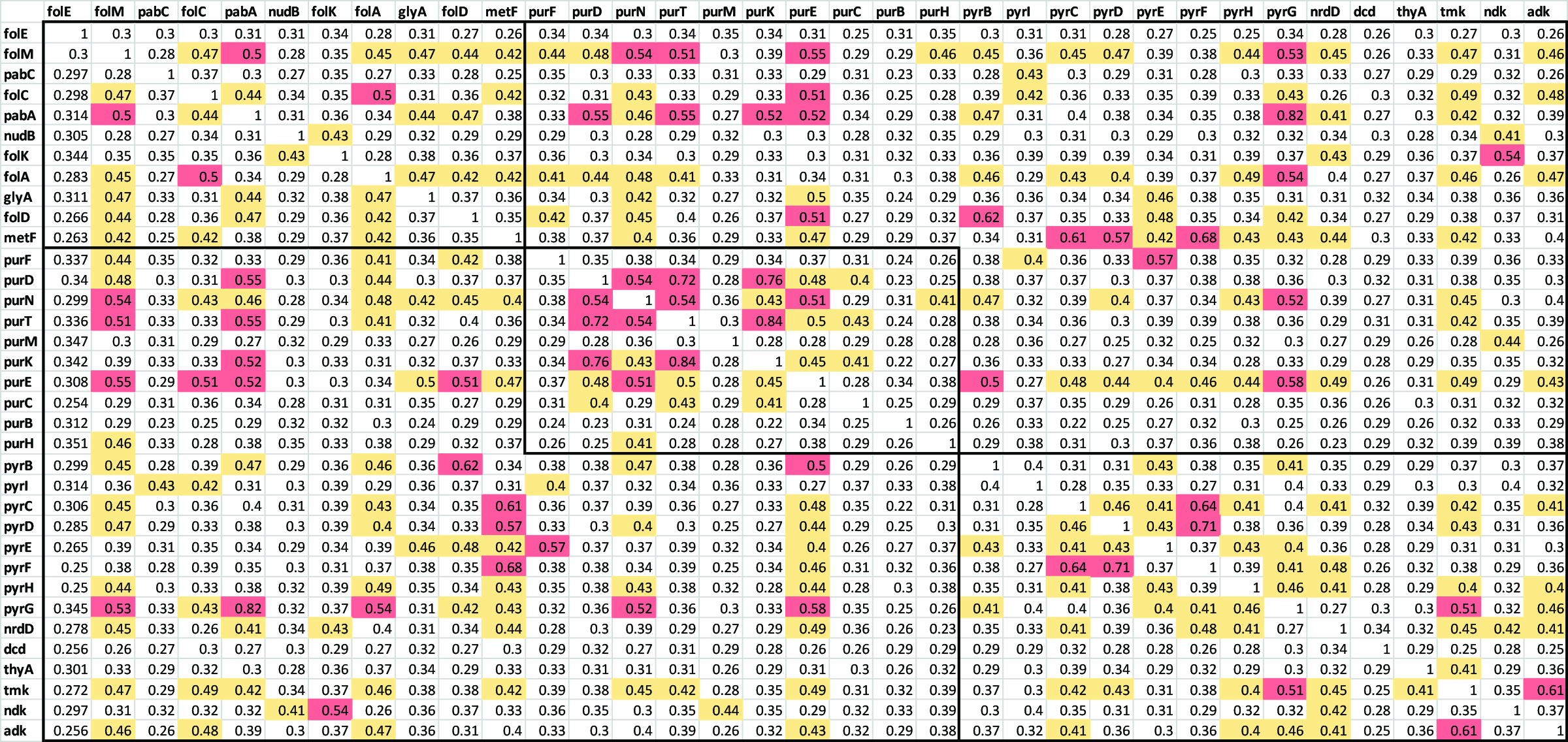

### Supplementary Fig 6

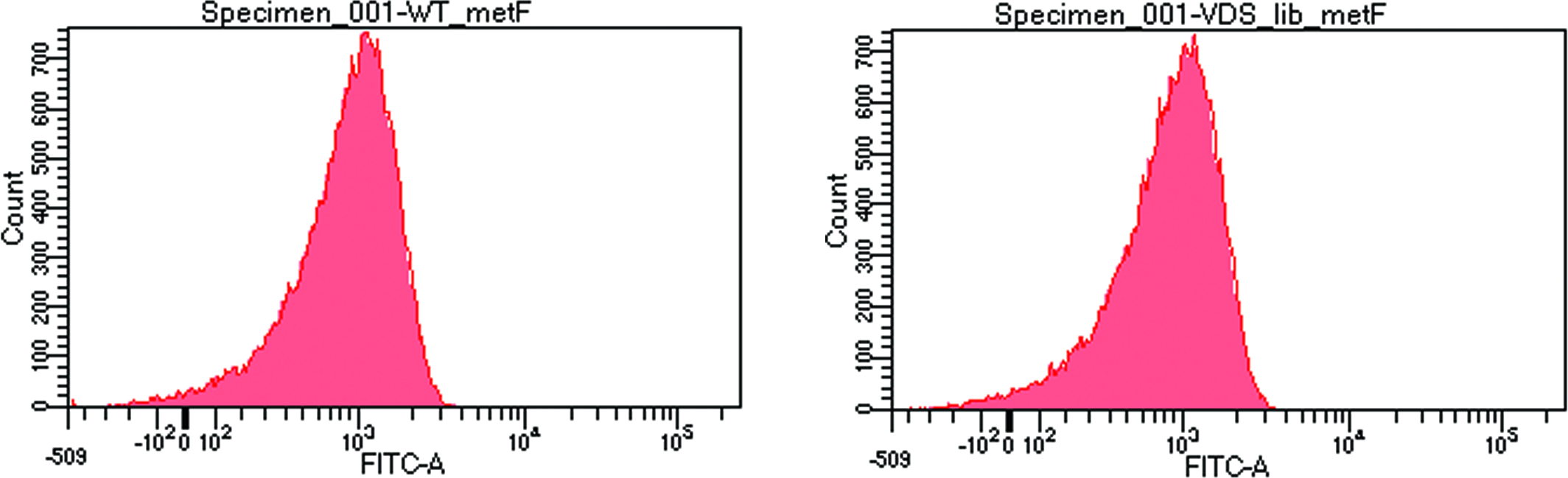

### Supplementary Fig 7

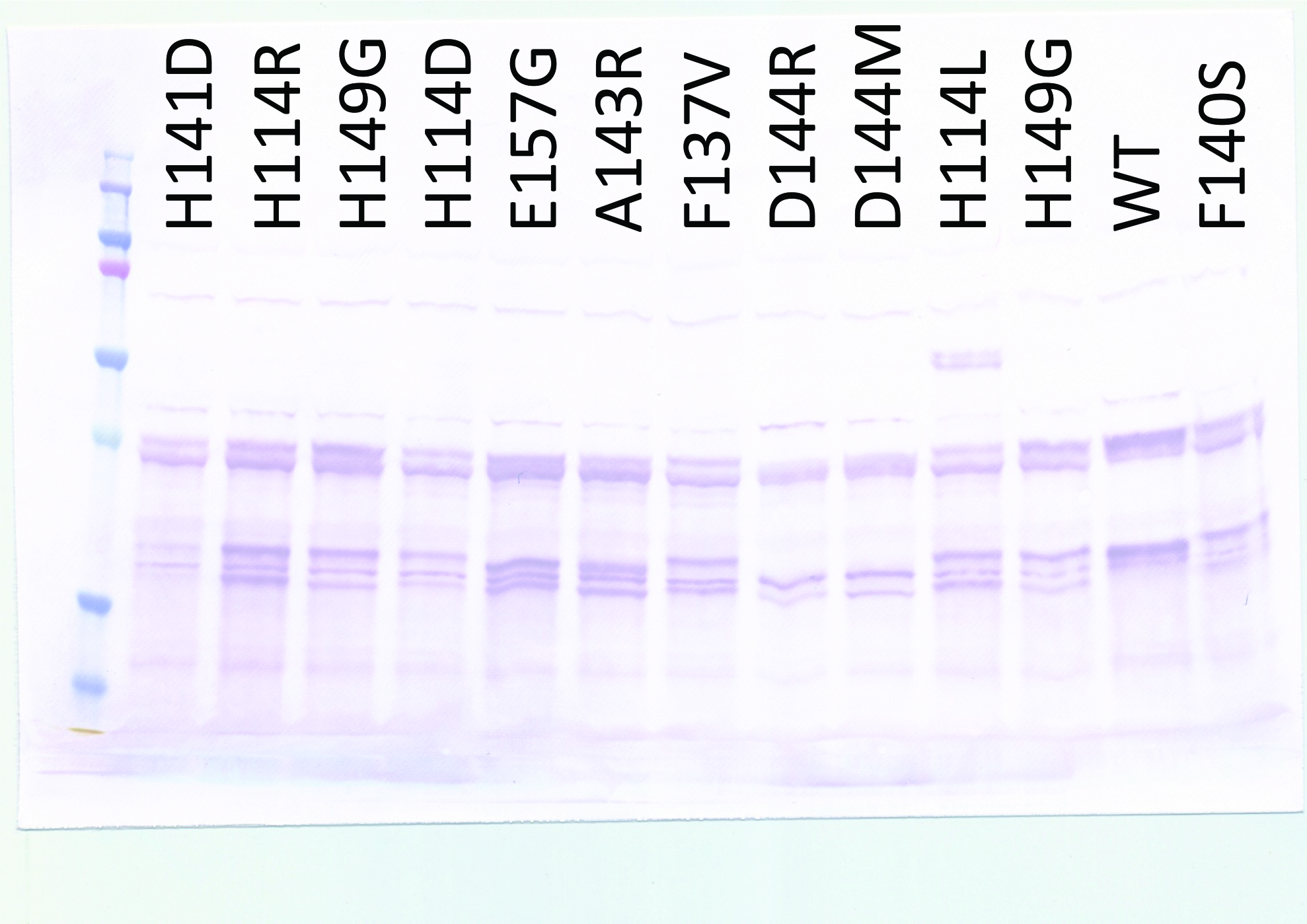

### Supplementary Fig 8

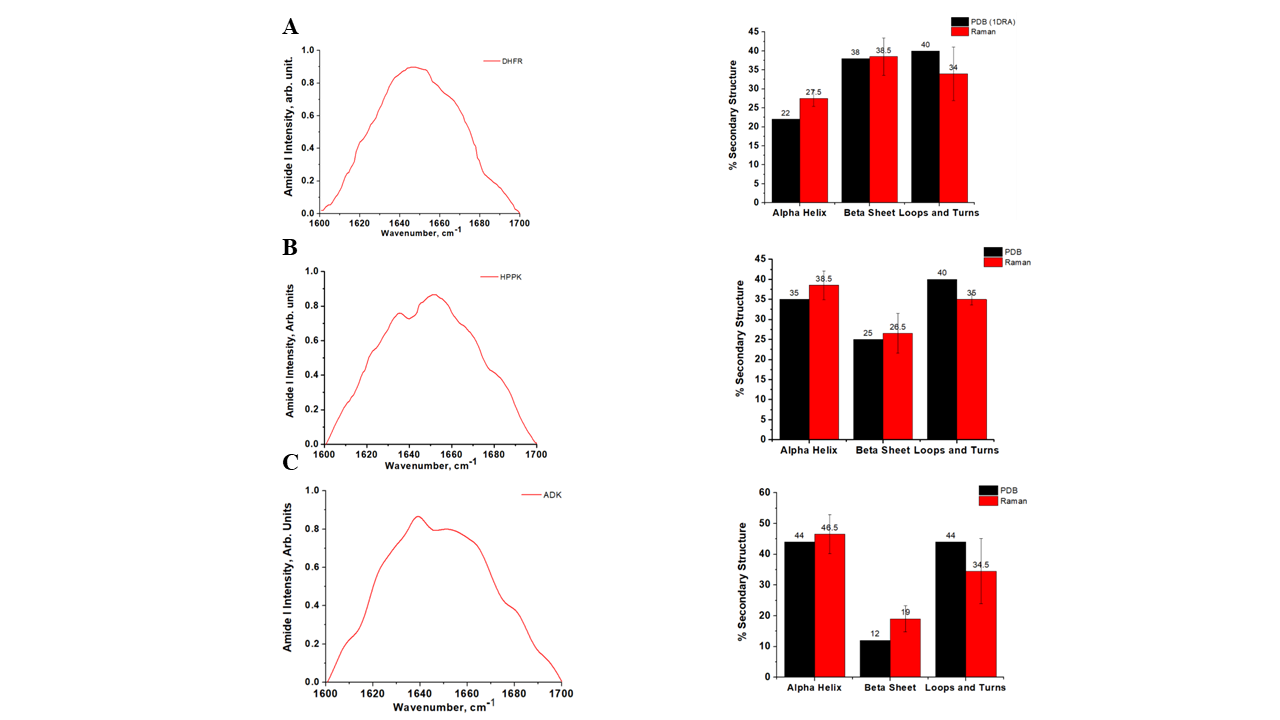

### Supplementary Fig 9

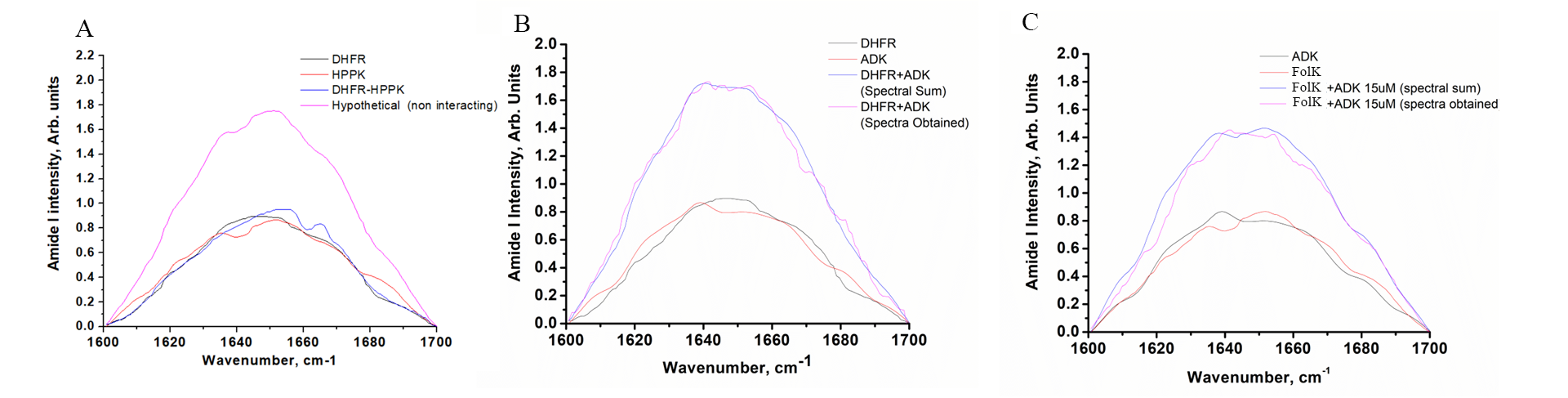

### Supplementary Fig 10

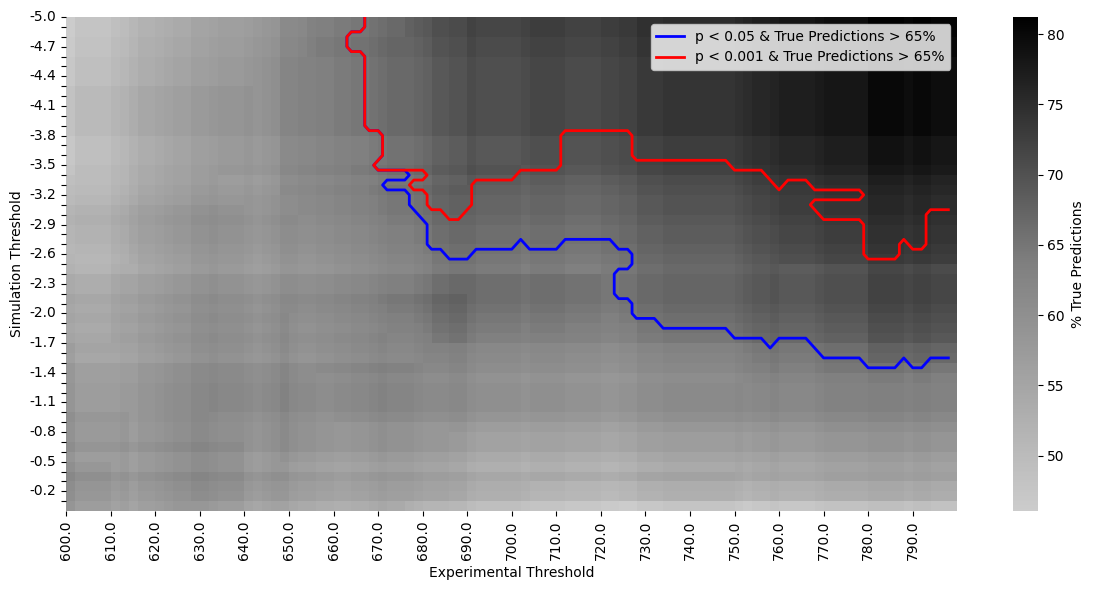

### Supplementary Fig 11

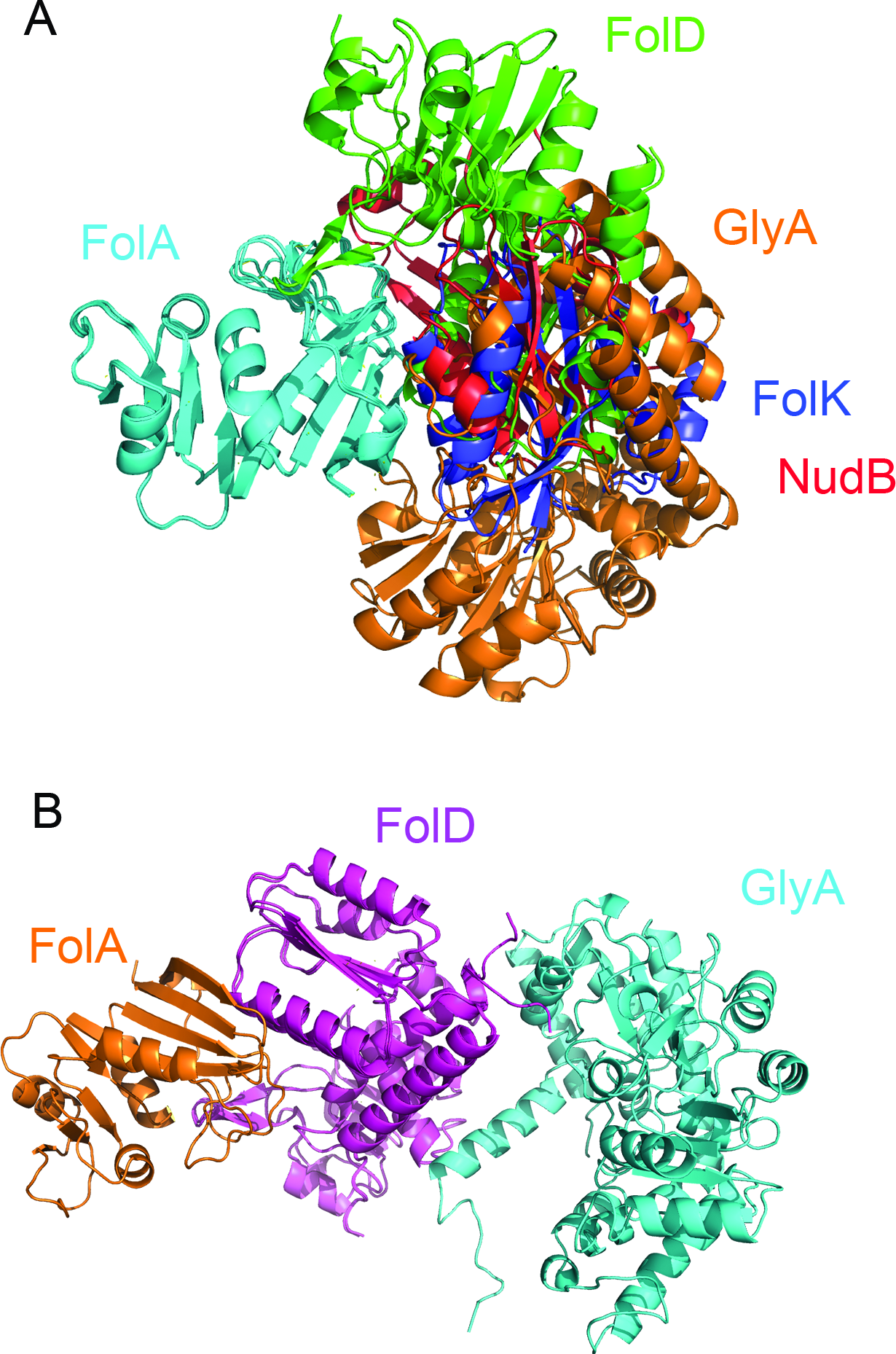

### Supplementary Fig 12

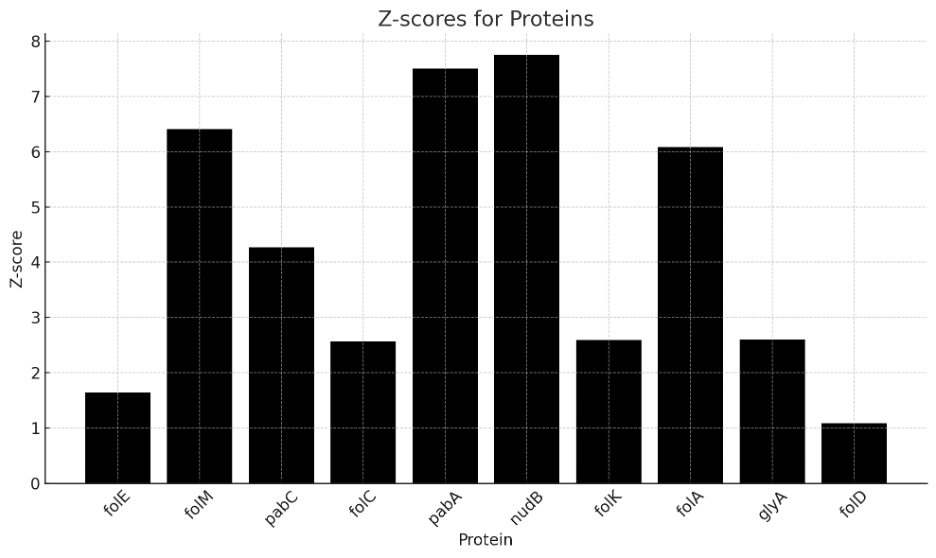

### Supplementary Fig 13

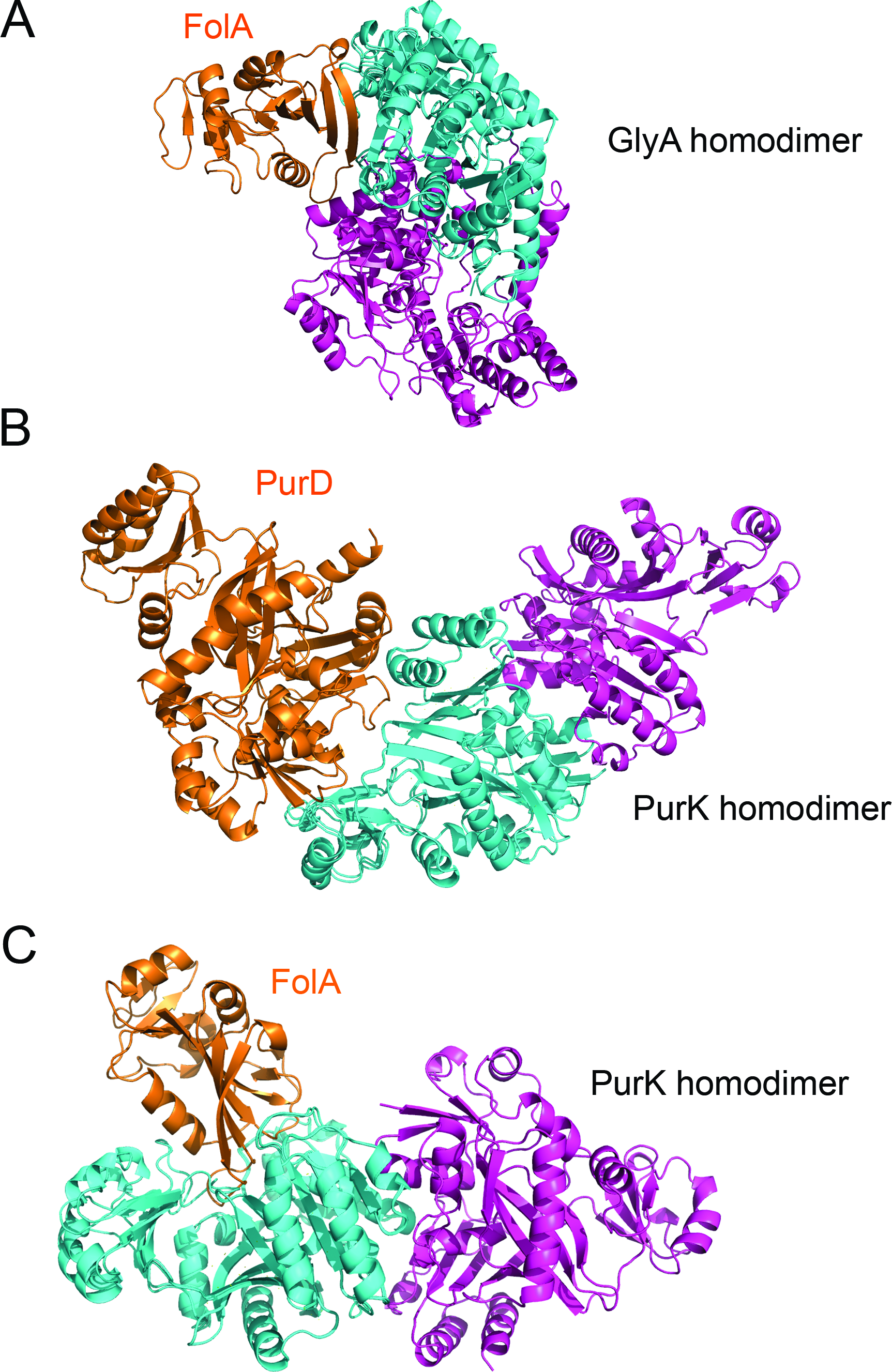

### Supplementary Fig 14

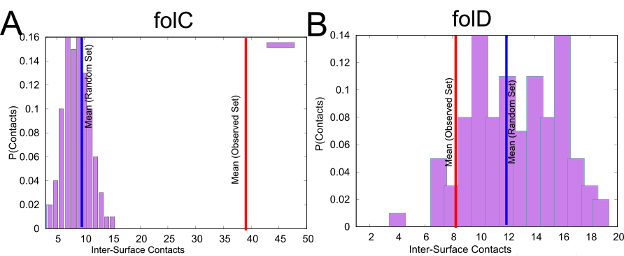

### Supplementary Fig 15

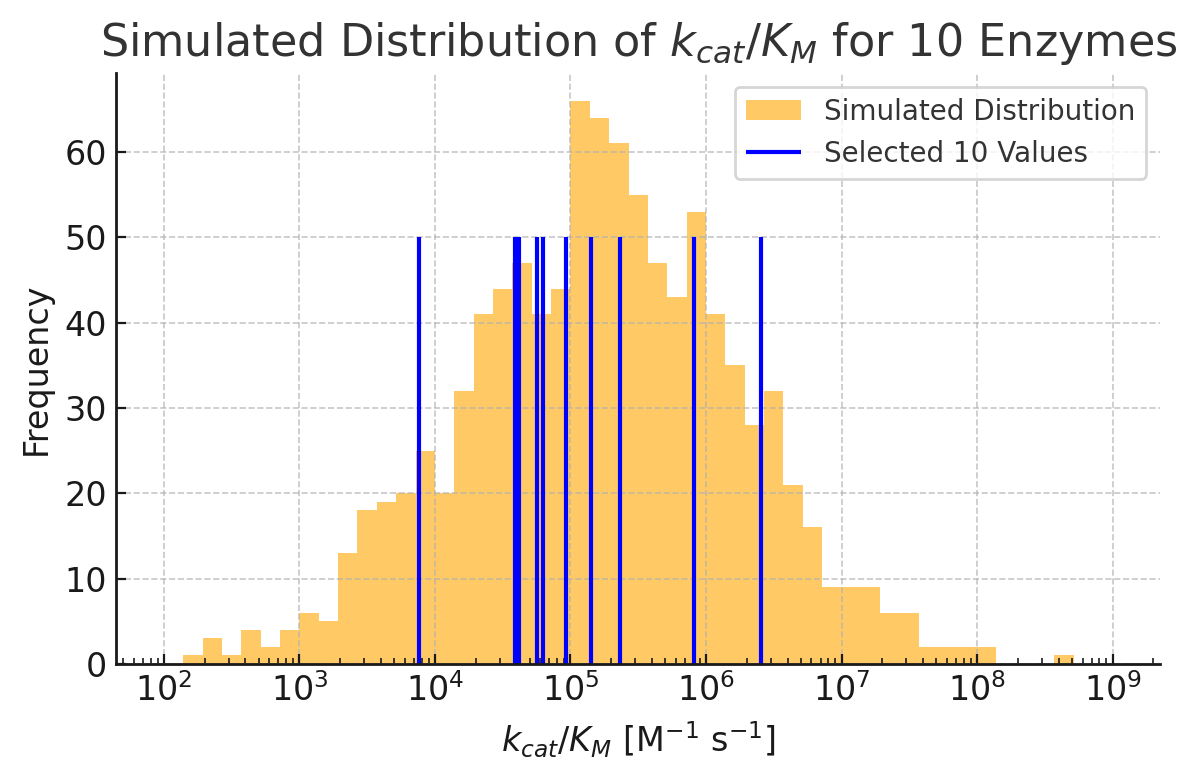

### Supplementary Fig 16

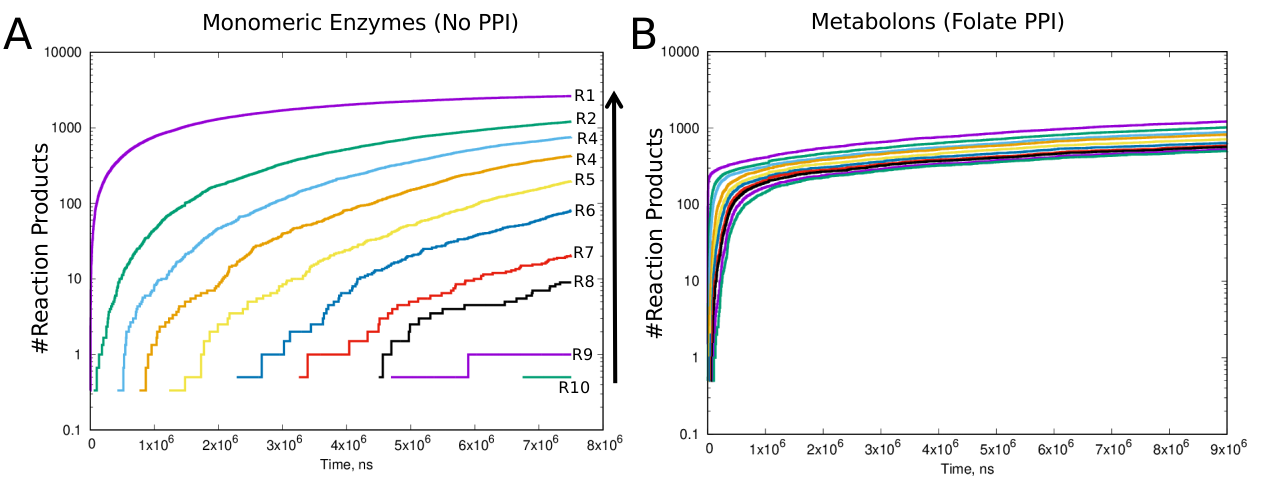

### Supplementary Fig 17

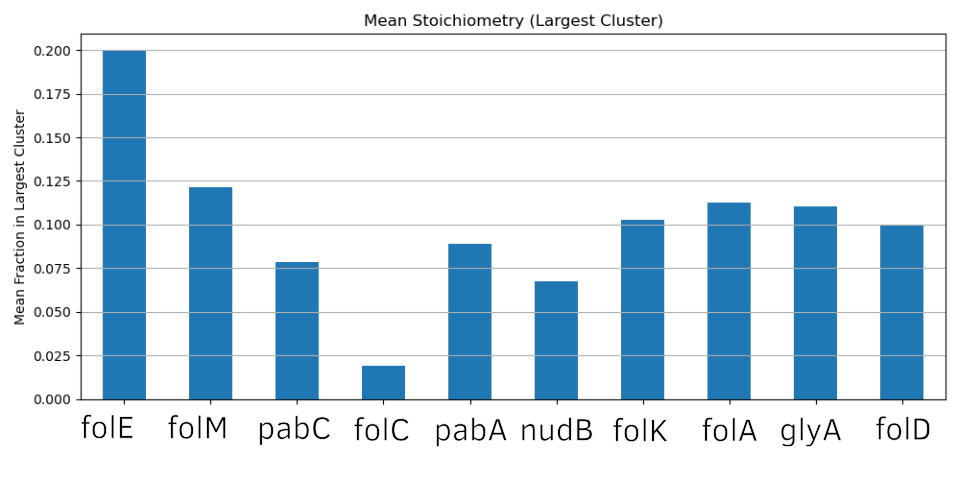
